# Structural insights into late-stage photosystem II assembly by Psb32

**DOI:** 10.1101/2025.05.08.652833

**Authors:** Stefan Bohn, Yat Kei Lo, Jan Lambertz, Jakob Meier-Credo, Torben Fürtges, Pasqual Liauw, Raphael Gasper-Schönenbrücher, Dennis Wiens, Julian D. Langer, Georg Hochberg, Eckhard Hofmann, Till Rudack, Marc M. Nowaczyk, Jan M. Schuller

## Abstract

Photosystem II (PSII) assembly is a stepwise process in which intermediate complexes with auxiliary proteins are transiently formed to allow efficient *de novo* biogenesis or repair of damaged PSII. In particular, the role of extrinsic PSII subunits (PsbO, PsbU, PsbV) and auxiliary proteins such as Psb27 for the formation and photoactivation of the Mn_4_O_5_Ca cluster, which catalyzes the unique water splitting reaction in mature PSII, remains unclear. Using cryo-electron microscopy, we have determined the structure of two novel late-stage PSII assembly intermediates from *Thermosynechococcus vestitus* BP-1. In contrast to previous studies, the resulting monomeric PSII complexes contain both PsbJ and Psb27 and exhibit a fully mature acceptor side, while the oxygen evolving complex (OEC) is still in an immature state. The second complex additionally associates with the late-acting assembly factor Psb32 and the extrinsic subunit PsbV. While Psb32 has received little attention, its proposed role in the complex challenges the previous assumption that all extrinsic subunits associate spontaneously, as well as the notion that PsbO initiates binding and solely drives OEC formation. Our structures of the Psb27-PSII and Psb32-PSII intermediates provide novel insights, how structural changes of C-termini of the D1 and D2 core proteins regulate maturation of the OEC and how the catalytic side is prepared for binding of the Mn_4_O_5_Ca cluster. The Psb32-PSII complex potentially represents the final PSII assembly intermediate that precedes the incorporation and photoactivation of the Mn_4_O_5_Ca cluster, allowing us to explain the final steps in the PSII biogenesis and assembly pipeline in great detail, as only the two extrinsic subunits PsbO and PsbU are missing.

## Introduction

Photosystem II (PSII) is a central membrane protein complex responsible for the light-driven oxidation of water in oxygenic photosynthesis^1^. In cyanobacteria, PSII exists as both a monomer and a dimer, with each monomer consisting of more than 20 subunits. The redox cofactors involved in intrinsic electron transport are mainly coordinated by the central proteins D1 (PsbA) and D2 (PsbD), which form the heterodimeric PSII core^2^. Surrounding them are the antenna proteins CP43 (PsbC) and CP47 (PsbB), which bind a large fraction of the chlorophyll *a* (Chl *a*) molecules responsible for absorbing light energy. This energy is then transferred to reaction center (RC) P680, where charge separation takes place. In addition, 13 low-molecular-weight proteins (LMWPs) bind to the complex and are located at the periphery, helping to bind cofactors and stabilize the complex^3^. Water oxidation is catalyzed by the oxy-gen-evolving complex (OEC), a Mn_4_O_5_Ca coordinated at the lumenal side of the complex^4,5^. In cyanobacteria, the extrinsic subunits PsbO, PsbU, and PsbV separate and protect the OEC from the thylakoid lumen^6^.

PSII biogenesis in cyanobacteria is subject to careful orchestration controlled by a number of assembly factors that do not occur in the functional complex^7,8^. Remarkably, the number of assembly factors involved in PSII biogenesis exceeds that of the subunits^9^. After formation of the reaction center complex (RCC), a pre-module composed of CP47 and several LMWPs associates, leading to the formation of RC47. This intermediate, together with the pre-existing CP43 module, forms a transient unit called PSII-I, which includes all major intrinsic subunits as well as the assembly factors Psb27, Psb28 and Psb34^10^. The presence of Psb28 and Psb34 leads to significant structural changes on the acceptor side of PSII. In particular, the altered orientation of D1-Glu244, D1-Tyr246 and D2-Glu241 prevents the association of bicarbonate (BIC) with the non-heme iron, similar to the reaction centers in non-oxygenic photosynthetic bacteria. This likely results in a shift of the Q_A_/Q_A_ redox potential towards a more positive value, protecting the immature complex from oxidative damage by facilitating safe charge recombination^10^. Psb27 binds to the lumenal side of the complex, where it stabilizes the E-loop of CP43 in a configuration similar to that of active PSII. This stabilization may support photoactivation of the OEC during PSII biogenesis^10–13^.

To date, all structurally resolved Psb28-containing complexes do not contain the LMWP PsbJ^10,14^, suggesting that PsbJ binding is the next step in PSII biogenesis after Psb28-PSII formation. It is hypothesized that PsbJ binding triggers structural reorganization leading to dissociation of Psb28 and Psb34^10,14^. Finally, the extrinsic subunits PsbO, PsbU, and PsbV bind to the lumenal PSII site and the OEC is formed in the process of photoactivation after the release of Psb27^7,15^. However, our knowledge of the exact sequence of events leading to the formation of active PSII remains limited.

Among the proteins thought to play a role in both the late steps of PSII assembly and repair is the protein TLP18.3/Psb32^16^. Psb32 abundance was found to be reduced in a PsbV deletion strain^17^, indicating a functional link between these two proteins and suggesting its potential role as a late-acting assembly factor. This protein family is conserved among plants, algae, and cyanobacteria^18^. The majority of the polypeptide contains a domain of unknown function (DUF477), also known as the TPM domain^19^. *In vitro* studies have shown that *A. thaliana* Psb32 (or AtTLP18.3) has low acid phosphatase activity^19,20^. Deletion mutants of Psb32 exhibit a distinct phenotype only under light stress conditions, characterized by impaired oxygen evolution and PSII repair, together with reduced levels of dimeric PSII^16^. However, the precise role of Psb32 in the PSII life cycle remains poorly understood.

In our current investigation, we have elucidated two previously uncharacterized structures of PSII intermediates using cryo-electron microscopy (cryo-EM). In contrast to previous studies focusing on immature PSII complexes, these complexes were isolated using Twin-Strep-tagged Psb27 without deletion of additional PSII components. The resulting monomeric PSII complexes contain both PsbJ and Psb27 and exhibit a fully mature acceptor side. In addition, one of these complexes is associated with PsbV and the late-acting assembly factor Psb32. This particular complex probably represents the ultimate PSII assembly intermediate that precedes incorporation of the Mn_4_O_5_Ca cluster and thus photoactivation.

## Results

Potential PSII intermediates with bound Psb27 were isolated by affinity chromatography using a Twin-Strep-tag (TS-tag) fused to the N-terminus of Psb27 (see Supplemental Fig. S1 and S2) from *Thermosynechococcus vestitus* BP-1 and subsequently analyzed by single-particle cryo-electron microscopy (see Supplemental Fig. S3A). Because these biogenesis intermediates are present at extremely low abundance, cryo-EM analysis required purification conditions that maximize stability and particle recovery. We therefore replaced DDM with lauryl maltose neopentyl glycol (LMNG), which forms exceptionally stable and uniform micelles that preserve native transmembrane architecture while remaining tightly associated with the protein^21^. This strong association allows extensive removal of free detergent without compromising integrity, enabling vitrification under detergent-minimized conditions that greatly increased particle yield. Using this method, we obtained two high-resolution maps, which allowed reliable modeling with excellent statistical parameters (see Supplementary Table S1 and Figs. S3B-D, S4). The first map, with a resolution of approximately 2.8 Å, termed Psb27-PSII, represents a monomeric PSII complex that includes nearly all membrane-bound subunits (including D1, D2, CP43, CP47, Cyt *b*_559_, PsbH, PsbI, PsbJ, PsbK, PsbL, PsbM, PsbT, PsbX, PsbZ, Psb30) in addition to the assembly factor Psb27. Notably absent are the extrinsic proteins of mature PSII (PsbO, PsbU, PsbV) and the small single transmembrane helix subunit PsbY (Fig. 1A). The second map, at 2.9 Å resolution, designated Psb32-PSII (see Fig. 1), also represents a monomeric assembly intermediate containing all proteins as in Psb27-PSII and additionally the extrinsic subunit PsbV and the assembly factor Psb32. The presence of all assigned proteins was confirmed by mass spectrometry analysis (see Supplementary Fig. S5, Supplementary Data 1 and 2). In addition, all cofactors characteristic of mature PSII are present in both assembly intermediates, with the exception of the Mn_4_O_5_Ca cluster.

**Figure 1:**
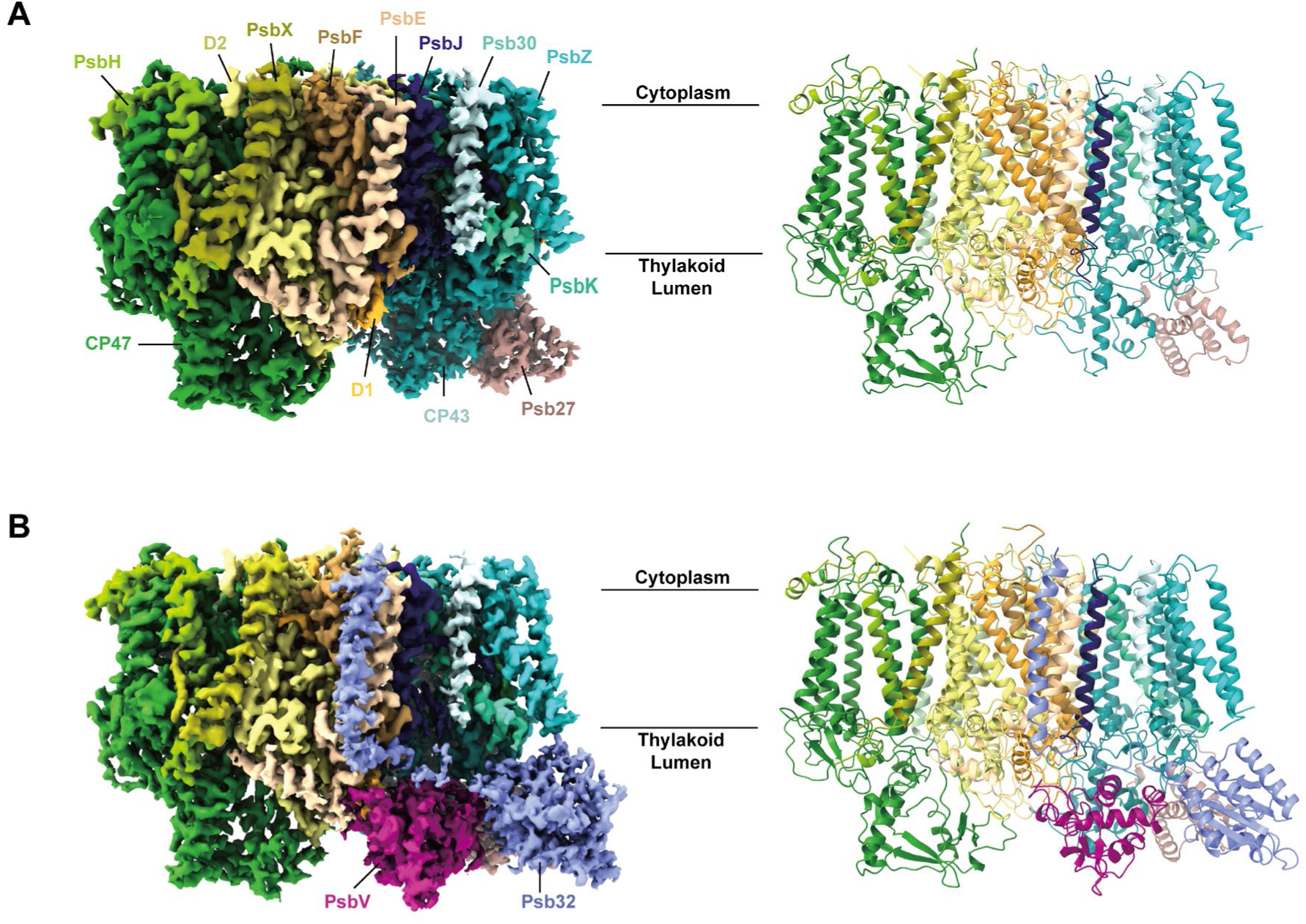
Structural overview of PSII assembly intermediates. Front view of the cryo-EM electron density and cartoon model of **A)** Psb27-PSII, and **B)** Psb32-PSII. Psb27-PSII is composed of the core PSII subunits: D1, D2, CP43, CP47, PsbE, PsbF, PsbH, PsbI, PsbJ, PsbK, PsbL, PsbM, PsbT, PsbX, PsbZ, Psb30, and the auxiliary protein Psb27. Psb32-PSII additionally binds Psb32 and the extrinsic subunit PsbV.

Previous structural studies of Psb27-containing PSII assembly intermediates were based on complexes that were either isolated from a *psbJ* knockout strain^10^ or PsbJ was potentially lost during sample preparation^14,22,23^, resulting in an empty Q_B_-binding pocket. Interestingly, Psb27-PSII and Psb32-PSII of this study both contain PsbJ and Q_B_ (Fig. 2). This suggests that the binding of PsbJ may complete the maturation of the PSII acceptor side by contributing to the formation of the Q_B_ binding pocket. The absence or low abundance of Psb28-containing PSII intermediates in our TS-Psb27 preparation from wild-type background suggests that PsbJ binding triggers or promotes the release of Psb28 as well as Psb34 from PSII-I to yield Psb27-PSII^10,14^.

**Figure 2:**
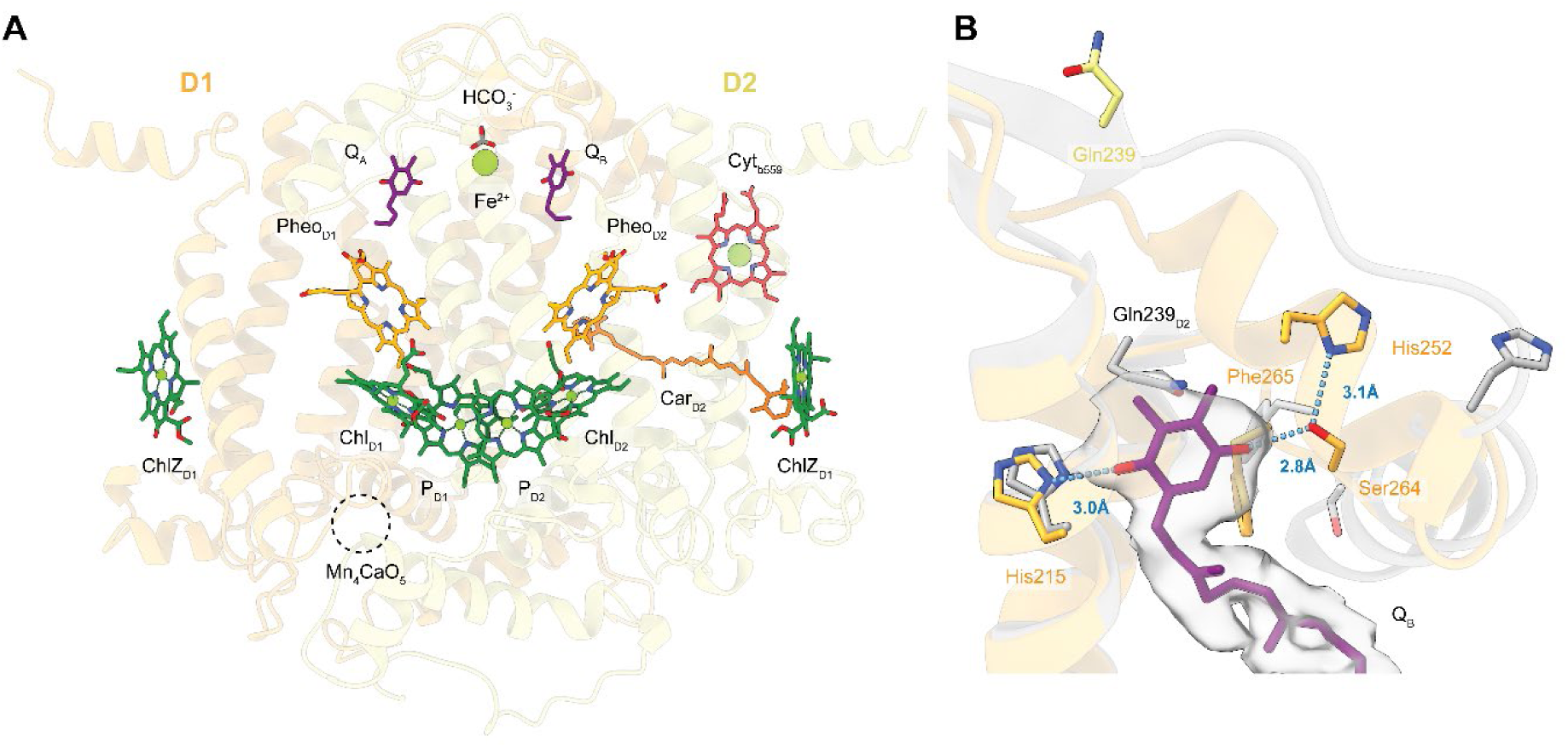
Cofactors of the Psb32-PSII assembly intermediate. **A)** Overview of cofactors assigned according to observed electron density in Psb32-PSII (PDB ID: 9I83). **B)** Zoom-in view of Psb32-PSII Q_B_ binding site, super-positioned with PSII-I (PDB ID: 7NHP) Psb32-PSII Q_B_ site adapted the mature PSII configuration, in which D1 His215, His252, and Ser264 are in position to coordinate Q_B_. D1 and D2 subunits of Psb32-PSII are colored in orange and yellow, respectively. PSII-I is colored in grey. The density of Q_B_ is shown at 5.8 σ.

To investigate the influence of Psb32-PSII complex formation on Psb32, we additionally resolved the structure of the soluble domain of Psb32 from *T. vestitus* fused to superfolder GFP (sfGFP) at a resolution of 2.12 Å by X-ray crystallography (see Supplementary Fig. S6, Supplementary Table S2). Our structural model (PDB-ID 8C7I) demonstrates an excellent overlap (Cα RMSD 1.74 Å, 149 atoms) with an earlier X-ray crystal structure of *A. thaliana* Psb32 (PDB-ID: 3PVH)^19^, except for the N-terminal helix 1, which shows partial unfolding in cyanobacterial Psb32 (see Supplementary Fig. S7).

There are no significant differences between the backbone fold of the soluble domain of PSII-bound Psb32 of the cryo-EM structure and the isolated domain of the X-ray crystal structure (Cα RMSD 0.66 Å, 149 atoms). However, Psb32 is a tail-anchored protein, and by a detailed examination of our Psb32-PSII cryo-EM density we identified its C-terminal transmembrane helix (TMH, residues D198 to S228), which is approximately 40 Å away from the soluble domain (Fig. 3A). The globular domain (residues S32 to K172) and the TMH are connected via a long linker (G173 to T197). The TMH is located at the position of PsbY in active PSII (PDB ID 4PJ0)^24^ near the heme of Cyt *b*_559_ (Fig. 3A). Psb32 and PsbY both interact with PsbE mainly due to hydrophobic interactions (Fig. S8). The proximity of the C-terminal helix of Psb32 to Cyt *b*_559_ is of particular interest as Cyt *b*_559_ is involved in alternative electron pathways within PSII that contribute to photoprotection^25^, consistent with the proposed role of Psb32 in protection against ROS^18^. Although the overall induced changes to the coordination of Cyt *b*_559_ are negligible within the limits of resolution, Psb32 could possibly affect its redox properties.

**Figure 3:**
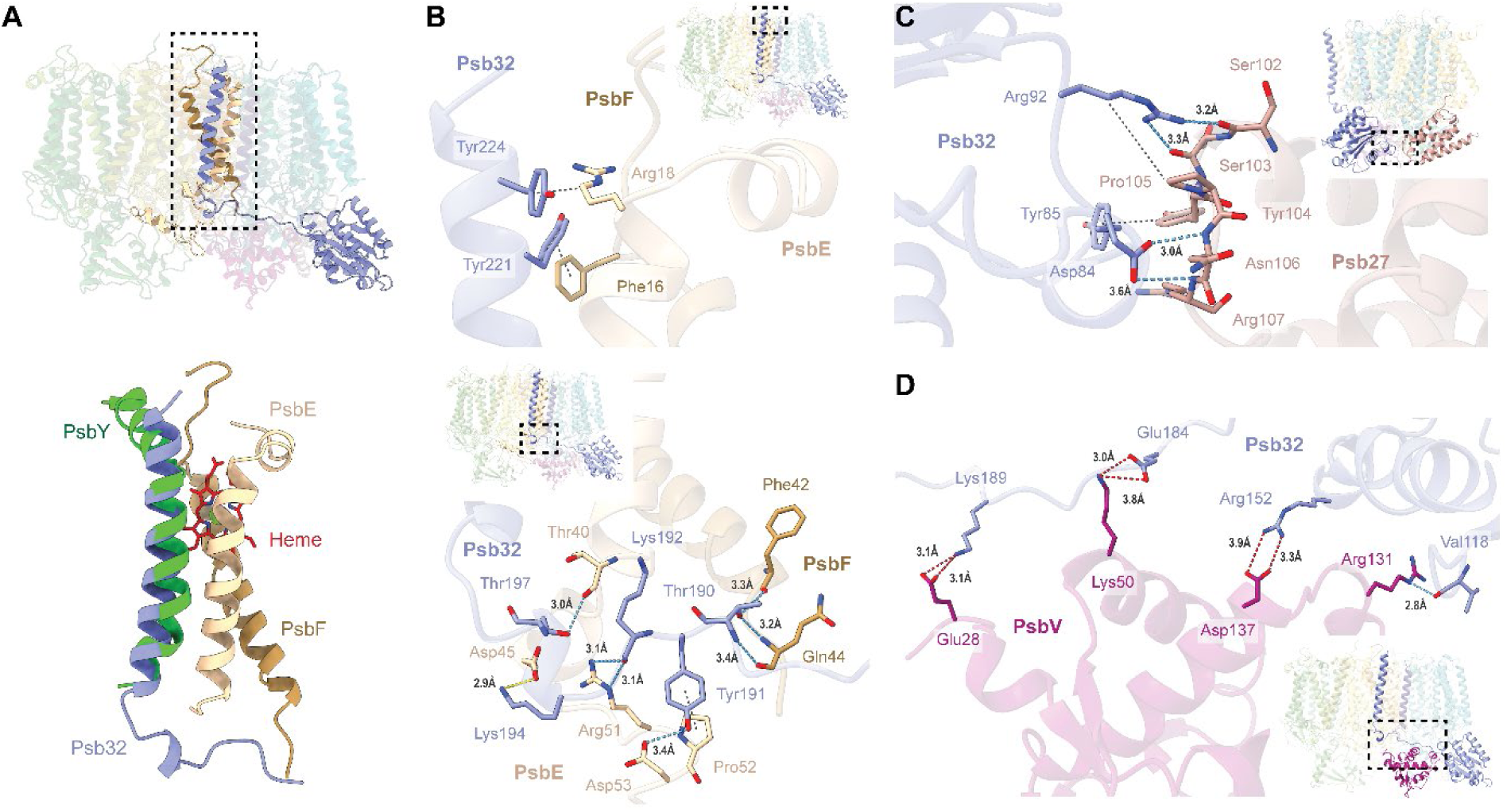
Psb32 interacts with PSII core subunits and assembly factors. **A)** Enlarged view of Psb32-PSII Cyt *b_559_* (boxed region), super-positioned with PsbY from the mature PSII (PDB ID: 4PJ0). In mature PSII, the heme group is enclosed by PsbE, PsbF, and PsbY, while in Psb32-PSII, the C-terminal transmembrane helix of Psb32 binds at the position of PsbY which can influence the potential of Cyt *b*_559_. **B-D)** Zoom-in view of specific interactions between Psb32, Psb27, PsbE, or PsbF with highlighted hydrogen bonds (blue dashed lines), salt-bridges (yellow dashed lines) and hydrophobic interaction pairs (grey dashed lines). Position of the interaction interface is illustrated in the inserts.

Only the Psb32 TMH interacts with the PSII core complex through PsbE and PsbF (Fig. 3B, C, Fig. S8, Table S3). The Psb32 soluble domain interacts with the assembly factor Psb27 and the extrinsic subunit PsbV (Fig. 3D, E). In the current structure, PsbV is located almost at its mature position near the lumenal loop regions of CP43 (residues 386-397 and 412-417) and D1 proteins (residues 305-315) (detailed in Supporting Table S3). The primary interaction motive between Psb32 and Psb27 is localized within the Psb32 loop from residues Leu80 to Pro89 and the Psb27 loop from residues Ser102 to Glu11. Precisely, the loop residues Arg82 and Asp84 of Psb32 as well as Ser102, Ser103, and Asn106, Arg107 of Psb27 form hydrogen bonds (Fig. 3D). Additionally, Psb32 Tyr85 and Psb27 Tyr104 form a specific van der Waals contact. The interface between Psb32 and PsbV is stabilized by salt bridges (Psb32 Arg152 with PsbV Asp137; Psb32 Glu184 with PsbV Lys50) and a hydrogen bond (Psb32 Val118 with PsbV Arg131) primarily involving the PsbV loop from residues Lys103 to Glu135 (Fig. 3E). Overall, these interfaces are relatively small, suggesting that the interaction between PsbV and Psb32 is transient and designed for easy detachment of Psb32 from the complex, a characteristic typical of biogenesis factors^26^.

Interestingly, while plastids encode only Psb32 (TLP18.3), cyanobacteria also encode a Psb32-like protein that differs from other bacterial Psb32-like proteins by possessing one or more long insertions near the C-terminus, which are not well conserved (Fig. S9). The precise function of this homolog and whether it could serve as an isoform capable of replacing Psb32 in the assembly process remains unclear. Structure predictions for the entire Psb32-PSII assembly intermediate with a hypothetical model replacing Psb32 with the Psb32-like protein, suggest that the putative transmembrane helix (TMH) of Psb32-like protein does not bind stably to PSII in contrast to a stable bound Psb32 THM (Fig. S10). The comparison of structure predictions of the isolated Psb32-like protein with Psb32 indicates a clear difference in prediction confidence and structure of the C-terminal tail (Fig. S11). Phylogenetic analysis shows that Psb32-homologs are found in a wide range of non-photosynthetic bacteria and that these homologs form an outgroup to the cyanobacterial proteins. Like the cyanobacterial variant, they carry a glycine-rich C-terminal extension, which is not present in Psb32 (Fig. S9). The loss of this extension thus may have played a role in the transition to chaperone functionality. Both Psb32-like proteins and Psb32 do not appear to be present in the basal branching cyanobacterial order *Gloeobacterales*. One possible explanation for this is that a Psb32-like precursor was transferred to an ancient cyanobacterium after *Gloeobacterales* branched off from other cynaobacteria and subsequently duplicated to give the Psb32 and Psb32-like clades in cyanobacteria. Alternatively, a Psb32-like protein existed in the last common ancestor of all cyanobacteria but was lost along the lineage to *Gloebacterales*. This protein would have then undergone duplication in the last common ancestor of all cyanobacteria excluding *Gloeobacterales*. These data show that Psb32 ultimately derives from a protein that originated in non-photosynthetic bacteria, which probably could not interact with PSII. Psb32 therefore most likely represents a case of neofunctionalization after gene duplication and a rare example in which the evolutionary origins of a novel assembly factor can still be reconstructed.

To further investigate the conformational changes required for OEC maturation, we compared the structures of PSII-I, Psb27-PSII, Psb32-PSII, and mature PSII in detail. Our analyses show no significant differences in the global fold between our biogenesis intermediates and the mature structure (Fig. S12A-D) but detailed changes in the OEC-coordinating residues. The most notable difference lies in the conformation of the D1 (His332-Ala344) and D2 C-terminal tails (Asn350-Leu352).

In the mature PSII structure, the D2 C-terminus points toward the D1 C-terminus (Fig. 4B), whereas in the immature Psb32-PSII structure, it is flipped away, obstructing PsbO binding. This observation aligns with previous studies that suggest reduced PsbO binding affinity in the presence of Psb27 and propose that simultaneous binding of these factors is transient^10,14,23,27,28^. In mature PSII, the D2 C-terminus interacts with both PsbU and PsbO (Fig 4C, D, Table S4). The spatial position of D2 Arg348 whose side chain forms a hydrogen bond with the PsbU Lys104 C-terminus of the mature conformation is maintained also in the immature conformation. However, a backbone-backbone interaction between D2 Ala351 and PsbU Gly101 is only possible in the mature conformation. The observed D2 terminus flip suggests that PsbU may stabilize the D2 C-terminus in a position that, in turn, could stabilize the D1 C-terminus. Whether the conformational change of the D2 C-terminus drives or results from the binding of the extrinsic subunits PsbO and PsbU is the focus of current ongoing studies.

**Figure 4:**
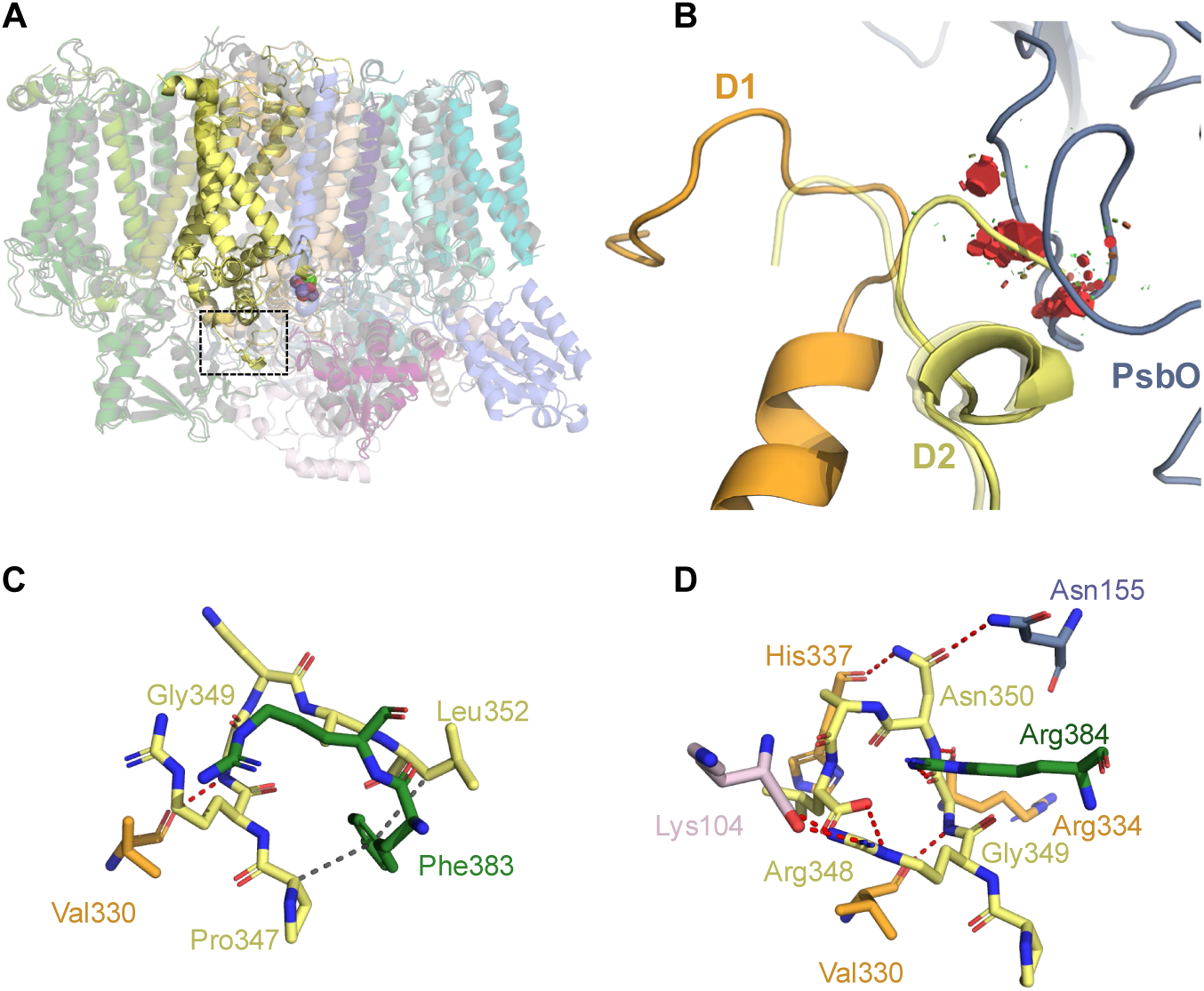
Conformational change of D2 C-terminal tail. **A)** Superposition of the mature PSII (grey) and Psb32-PSII complex (subunit coloring as in Fig. 1). **B)** Close-up of the D2 terminal region (as marked in A) reveals a steric clash between the D2 C-terminal tail residues in Psb32-PSII and PsbO in the mature PSII. In contrast the D2 C-terminus of the mature PSII (transparent) is position towards the D1 C-Terminus. **C)** D2 C-terminus stabilizing interactions in Psb32-PSII. The D2 C-terminus is mainly stabilized by CP47 (green). **D)** D2 C-terminus stabilizing interactions in mature PSII. The terminus interacts with D1 (orange), PsbU (pink), and PsbO (blue).

In all our structures at least the end of the C-terminal tail of D1, which contains the residues D1-Asp342 and D1-Ala344 involved in coordination of the Mn_4_O_5_Ca cluster, is highly flexible, as observed in other PSII biogenesis intermediates earlier. Consistent with this, no clear electron density corresponding to the Mn_4_O_5_Ca cluster is observed, though some weak density at low thresholds indicates minor sub-stoichiometric binding or the presence of mixed populations that cannot be separated by further classification (Fig. S13D-F). This concludes that the observed Psb32-PSII intermediate is in an immature state, in which the residues involved in binding of the Mn_4_O_5_Ca cluster are already in their final position as in mature PSII – except for D1-Asp342 and D1-Ala344 - but the Mn_4_O_5_Ca cluster itself is mainly not present or not yet stably bound. A small subpopulation of particles may contain a fully assembled Mn_4_O_5_Ca cluster, but slightly displaced from its mature position, as indicated by a weak density in this area. The observed displacement could either mean that the cluster is stalled in a non-functional state, or it could also be an artifact of radiation damage. All in all, binding of PsbO (and PsbU) appears necessary to finally position D1 Asp342 and Ala344 in a catalytic active position, enabling the formation, photoactivation and stable binding of the Mn_4_O_5_Ca cluster to achieve final maturation of the OEC-coordinating residues.

We identified D1 His332 and D1 Glu333 as key residues reflecting the progression of the OEC towards its mature state. In the PSII-I structure, His332 adopts an immature conformation, oriented away from the Mn_4_O_5_Ca cluster binding site and coordinating a chloride ion via its protonated Nδ atom (Fig. 5A). In Psb27-PSII, His332 exhibits a mixture of both the immature, chloride-coordinating conformation and the mature conformation, in which D1 His332 interacts with D1 Glu329 (Fig. 5B). In contrast, in Psb32-PSII, D1 His332 shifts to its mature orientation, enabling coordination of the Mn_4_O_5_Ca cluster. In both the mature PSII and Psb32-PSII structures, the mature conformation of D1 His332 is stabilized by a hydrogen bond between the backbone of D1 Glu329 and the protonated D1 His332 Nδ atom (Fig 5C). Similarly, D1 Glu333 undergoes a pronounced conformational change between Psb27-PSII and Psb32-PSII. In Psb27-PSII, the backbone position of D1 Glu333 suggests its side chain is oriented away from the Mn_4_O_5_Ca cluster binding pocket similar as in PSII-I (Fig. 5D). However, in Psb32-PSII, the density for D1 Glu333 reveals a side chain shift enabling to point towards the Mn_4_O_5_Ca cluster, adopting a mature conformation (Fig. 5E). All other resolved residues that coordinate the Mn_4_O_5_Ca cluster in the mature complex, namely D1 D170 as well as CP43 E342 and R345, have already adopted their mature conformation, prepared for OEC incorporation (Fig. 5F). Our structural analysis indicates that the con-formational transitions of His332 and Glu333 act as a critical molecular switch, governing the assembly of the OEC. Notably, Psb32 and PsbV appear to initiate these transitions, underscoring their role during the final steps of PSII biogenesis.

**Figure 5:**
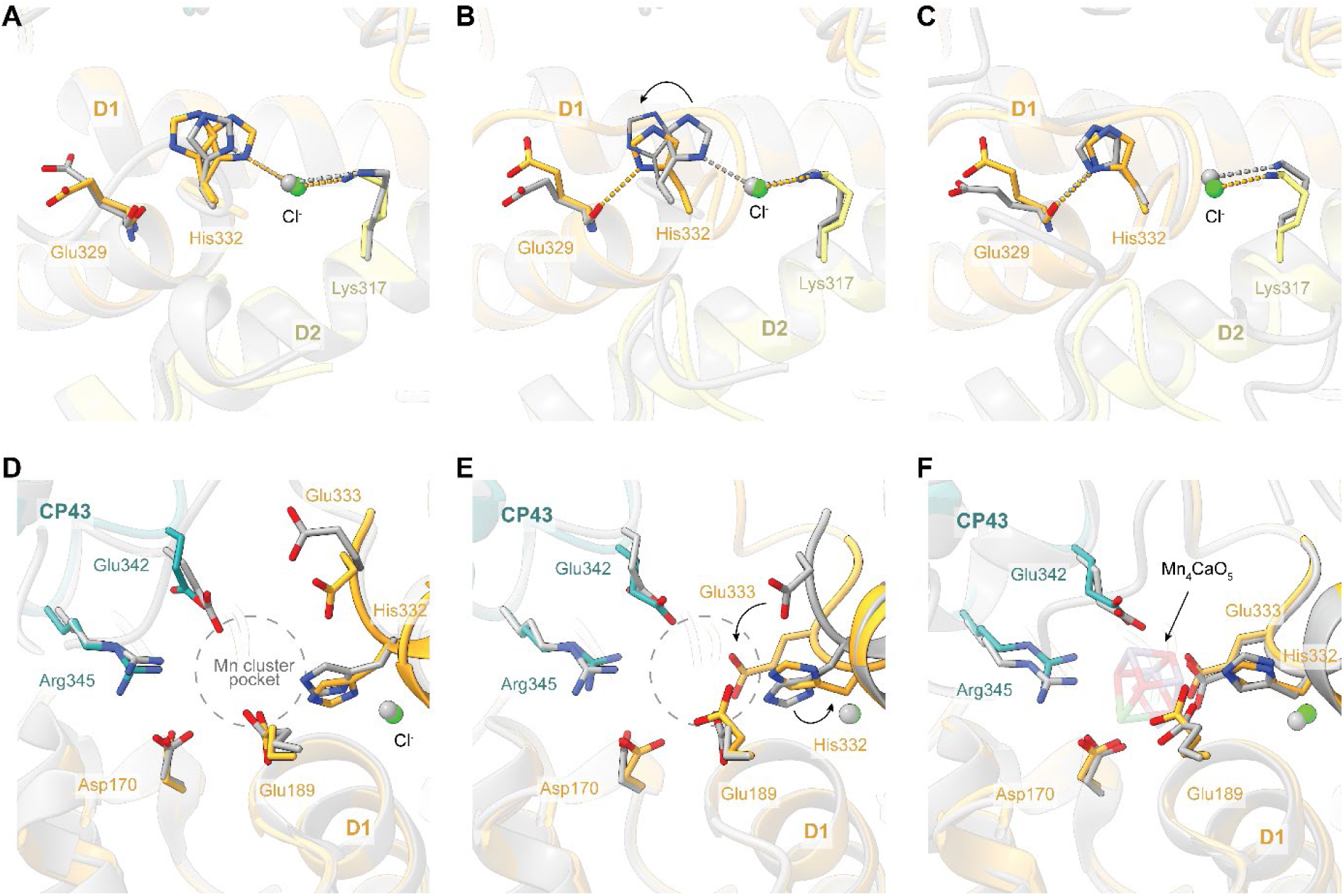
Maturation of the Mn_4_O_5_Ca pocket. **A)** Superposition of PSII-I (grey, PDB-ID: 7NHP) and Psb27-PSII (PDB-ID 9I82), **B)** Psb27-PSII (grey) and Psb32-PSII (PDB-ID 9I83), and **C)** Psb32-PSII and mature PSII (grey, PDB-ID: 4PJ0). In **A-C** the conformational changes of His332 of D1 are highlighted. **D)** Superposition of OEC between PSII-I (grey) and Psb27-PSII, **E)** Psb27-PSII (grey) and Psb32-PSII, and **F)** Psb32-PSII and mature PSII (grey). In **D-F** residues that coordinate the Mn_4_O_5_Ca cluster in the mature state are highlighted. In **B** and **E**, arrows indicate major conformational changes between Psb27-PSII and Psb32-PSII. Density fitting of the shown residues is illustrated in Fig. S13.

## Discussion

In our study we were able to elucidate the structure and function of the Psb32-PSII assembly intermediate. Our findings indicate that the assembly factor Psb32 interacts with the extrinsic subunit PsbV and with Psb27, a prominent factor previously described to facilitate the attachment of the CP43 antenna module to RC47^10,13,14^. The tail-anchored protein Psb32 is comprised of two major functional parts, and we hypothesize that they fulfill distinct roles within the assembly complex.

The C-terminal transmembrane helix of Psb32 binds near Cyt *b*_559_ and occupies the same position as PsbY in the mature complex. Although the overall induced changes to the coordination of Cyt *b*_559_ are neglectable within the error of resolution, Psb32 may influence its redox properties. This is of im-portance because Cyt *b*_559_ can switch between four redox forms (high potential: Em of +370–400 mV; intermediate potential: Em + 170–250 mV; low potential: Em + 70–100 mV; very low potential: Em−45 mV) but the structural origin for this redox-switching mechanism is still unknown^29^. The high potential (HP) form is characterized by an at least temporarily reduced heme iron, which can’t be oxidized by oxygen. Therefore, it can serve as an alternative electron donor for P680+. Instead, all low potential forms of Cyt *b*_559_ can be easily oxidized by oxygen and thus have the capability to accept an electron from the PSII acceptor side. By switching between the HP and LP form, Cyt *b*_559_ is involved in photo-protection by providing cyclic electron flow between the PSII acceptor and donor side^30,31^. This protection mechanism operates in PSII complexes without a functional OEC and has been shown to be important for its photoactivation^32^. Moreover, Cyt *b*_559_ of PSII without PsbY is also locked in the low potential state but little is known about the mechanism by which it switches between the two states^33^.

The lumenal domain of Psb32 is mainly coordinated by PsbV and Psb27 and may facilitate its attachment to the PSII complex. This is an unexpected finding, since the last steps, the association of the extrinsic subunits, were thought to be a spontaneous and unassisted process based on *in vitro* experiments^34^. In the cell, the situation may be different, and Psb32 can control the direction of the process and improve the efficiency of the assembly process. Based on our structure, a possible scenario might be that Psb32 stabilizes the binding of PsbV, which in turn is necessary to recruit PsbO to the complex in the next step to flip the D2 C-terminus into the mature position. This would explain why PsbO is not bound to any intermediate complex until the Mn_4_O_5_Ca cluster is integrated.

Furthermore, the above considerations for the role of Psb32 in PSII assembly are also consistent with its predicted role within the repair cycle^18^, as the reassembly of damaged complexes overlaps with the stages of *de novo* biogenesis described above^7^. Psb32 was shown to be associated with active PSII, protecting the complex from peroxide^18,35,36^ which may be related to its influence on Cyt *b*_559_ in Psb32-PSII. Besides the protection mechanism, it might be possible that binding of Psb27 is increased in presence of Psb32, initiating repair by anchoring Psb27. This suggests that Psb32 may play a role in recruiting Psb27, in addition to its involvement in assembly. This could provide an alternative explanation for the reduced PSII recovery rates observed in a Psb32 deletion strain following photoinhibition by high light^18^.

By integrating these new findings, we can now extend the existing model of the PSII assembly (Figure 6). Following the formation of the reaction center complex (RCC), a pre-module consisting of CP47 and several low molecular weight proteins (LMWPs) attaches, resulting in the formation of RC47. This complex is stabilized by the action of Psb28 and Psb34 that induces significant structural alterations on the acceptor side of PSII that prevent binding of bicarbonate (BIC) to the non-heme iron. This may shift the Q_A_/Q_A_- redox potential to a more positive value to promote safe charge recombination^10,37^, which protects the immature complex from oxidative damage. Subsequently, RC47 associates with the CP43 module. Previous work demonstrated that Psb27 is already bound to free CP43, where it might protect free CP43 from degradation or maintaining the E-loop in a conformation suitable for its subsequent association with the RC47 complex^13,27^. The structure of the resulting assembly intermediate revealed how a not-yet-completely assembled but already partially active molecular machine is protected from photodamage^10,14^. In this intermediate the Q_B_ site is not matured. However, our new structures show that following the dissociation of Psb28 and Psb34 and the binding of PsbJ, the Q_B_ site matures, with a molecule of PQ bound, similar to mature PSII. This maturation is a prerequisite for forward electron flow^15^. Apart from the binding of extrinsic subunits, this complex now seems to be intact. However, there are still side chains around the OEC that are not yet in a mature conformation, verifying that Psb27 on its own is not sufficient to promote photoactivation. Upon maturation of the acceptor side, Psb32 binds to the complex and, interestingly, the coordination site progresses towards maturation, as indicated by the conformational shifts in His332 and Glu333. These key residues are now in their final position to coordinate the Mn_4_O_5_Ca cluster, suggesting that Psb32, along with PsbV, plays a crucial role in initiating these transitions. All in all, the structural transitions underline the role of both Psb32 and PsbV in the final steps of PSII assembly. Furthermore, the critical role of PsbO in photoactivation becomes increasingly clear, though its action relies on a well-prepared environment orchestrated by the other biogenesis factors. Following photoactivation, Psb32 is replaced by PsbY, a shift that promotes forward electron transfer over cyclic electron transfer via Cyt *b*_559_ that offered protection against photoinhibition in the yet inactive complex.

**Figure 6:**
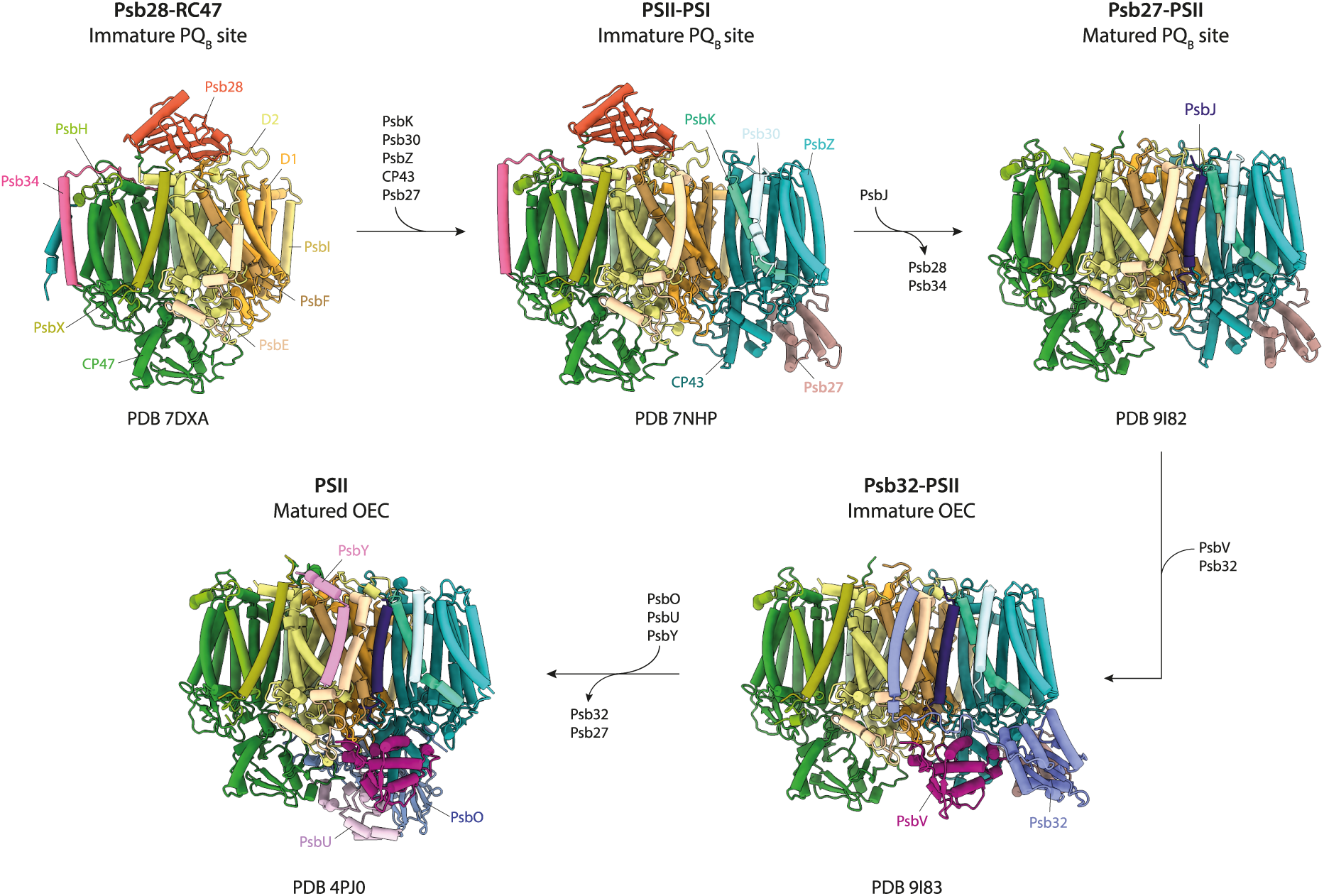
Schematic model of the late PSII assembly process after formation of Psb28-RC47. Association of the CP43 module to RC47 is stabilized by the assembly factor Psb27. Upon binding of the LMWP PsbJ, Psb28 and Psb34 are released from Psb27-PSII, forming the matured PQ_B_ site. Psb32 aids the assembly of the extrinsic subunit PsbV. Additionally, Psb32 prevents binding of PsbY, thus possibly protecting the immature intermediate from redox damage. Maturation of PSII is followed by association of the remaining subunits, as well as release of Psb32 and Psb27.

With this research, we now have a comprehensive understanding of the final steps in the PSII biogenesis and assembly pipeline, with only PsbY and the extrinsic subunits PsbO and PsbU remaining to be incorporated.

## Material and methods

### Mutant generation and cultivation

A *Thermosynechococcus vestitus* BP-1 mutant line was created by modifying the N-terminal sequence of Psb27 with a Twin-Strep-Tag after the N-terminal Cysteine (Sequence: C_mod_ANV-GGGSSAW-SHPQFEKGGGGSSGGGSAWSHPQFEKGGGS-Psb27). The DNA-sequence was synthesized by Twist Bioscience (USA) and transferred into the pJET1.2 vector with the CloneJET PRC-cloning kit according to the manufacturer (Thermo Fisher Scientific). A restriction site was integrated 170 bp upstream of Psb27, in which a kanamycin resistance cassette was integrated by ligation. Segregation was confirmed by PCR using forward (GAGCGATATCCATCCACGGTTGCCAAGAG) and reverse (GCCGAATTCTACCCTAACCCGCTGCTGTC) primers binding before and after the resistance cassette, respectively. The strain was grown in 25-L photobioreactors (Bioengineering) in Bg-11 media using in presence of 150 µg·ml^-1^ kanamycin^28^. The cells were harvested, pelleted and the membranes extracted with a parr bomb as described previously^28^.

### Protein purification

PSII complexes were extracted from thylakoid membranes and purified by Strep-tag affinity chromatography as described previously using Strep-Tactin®XT® columns (IBA Lifesciences)^38^. The final buffer of the protein complexes was PSII-DDM-buffer (20 mM MES pH 6.5, 10 mM CaCl_2_, 10 mM MgCl_2_, 500 mM mannitol with 0.03% (w/v) n-dodecyl-ß-d-maltoside (DDM)).

### Detergent-exchange and removal

In scope of the cryo-EM sample preparation we exchanged the detergent DDM to lauryl maltose neopentyl glycol (LMNG) directly on Strep-Tactin®XT® columns (CV = 5 ml). The sample was applied to the column in PSII-DDM-buffer, which was was exchanged to PSII-LMNG-buffer (PSII-buffer with 0.05% (w/v) LMNG) in a gradient from 20% to 100% over 8 CV. After full removal of the PSII-DDM-buffer (after 1 CV 100% PSII-LMNG-buffer), a second gradient from 80% to 0% PSII-LMNG-buffer against a detergent-free PSII-buffer was done over 6 CV. Proteins were eluted with 50 mM biotin in PSII-buffer without detergent after 1 CV of washing at 100% PSII-buffer without detergent. During all steps, a flow rate of 1 ml·min^-1^ was applied.

### Biochemical characterization

Oxygen evolution measurements were done as described previously^28^ with isolated protein equal to 2-5 µg·ml^-1^ Chl in presence of 1 mM 2,6-Dichloro-1,4-benzoquinone (DCBQ) and 5 mM ferricyanide. Blue-native PAGE and urea-based SDS-PAGE were done as described previously^39,40^. As size standard, Thermofisher PageRuler Prestained protein ladder #26616 or purified PSII^38^ were applied.

### MALDI-ToF MS

MALDI-ToF MS measurements were performed as described by^38^. Briefly, the target subunits were purified using C18-ZipTips (Merck, Germany). The eluted organic fractions were lyophilized in a vacuum concentrator (Eppendorf, Germany), reconstituted in 0.1% Trifluoroacetic acid (TFA) and mixed in a 1:1 ratio with sDHB (Bruker Daltonics, Germany) matrix solution (50 mg/ml in 50% (v/v) Acetonitrile (ACN), 50% (v/v) water and 0.1% (v/v) TFA). Subsequently 1 μl aliquots of the mixture were deposited on a BigAnchor MALDI target (Bruker Daltonics, Germany) and allowed to dry and crystallize at ambient conditions.

MS spectra were acquired on a rapifleX MALDI-TOF/TOF (Bruker Daltonics, Germany) in the mass range from 3000-7000 m/z in reflector positive mode. The Compass 2.0 (Bruker, Germany) software suite was used for spectra acquisition and processing.

### LC-MS

Protein samples were reduced with TCEP and cysteines alkylated with IAA (Thermo Fisher, Germany). Subsequent proteolytic digests were performed using S-TRAPs (Protifi, USA) according to the manufacturer’s instructions. Peptides were further desalted and purified on Isolute C18 SPE cartridges (Biotage, Sweden) and dried in an Eppendorf concentrator (Eppendorf, Germany) as described by^10^. After solubilization in 0.1% (v/v) formic acid (FA) in ACN/water (95/5, v/v), samples were subjected to LC-MS/MS analysis on a nanoElute (Bruker, Germany) system, equipped with C18 analytical column (15 cm * 75 µm, particle size: 1.9 µm (PepSep, Denmark)) coupled to a timsTOF Pro 2 mass spectrometer (Bruker, Germany). Samples were loaded directly onto the analytical column with twice the sample pick-up volume with buffer A. Peptides were separated on the analytical column at a flow rate of 500 nl/min with the following gradient: 2 to 35% B in 17.8 min, 35 to 95% B in 0.5 min and constant 90 % B for 2.4 min with buffer A (0.1% FA in water) and buffer B (0.1% FA in acetonitrile). Peptides eluting from the column were ionized online using a captive spray ion-source and analyzed in DDA-PASEF mode with a cycle time of 100 ms and 4 PASEF-MS/MS events. Spectra were acquired over the mass range from 100-1700 m/z and a mobility window from 0.85-1.3 Vs/cm^2^.

Data analysis was performed in Fragpipe 17.1 using MSFragger 3.4 for database searches^41^. Raw files were recalibrated and searched against the Uniprot proteome for *Thermosynechococcus vestitus* (UP000000440; obtained 2022-03-17) along with the modified Psb27 sequence. The search space was restricted to tryptic peptides with a length of 7-50 amino acids, allowing for up to two missed cleavages and with a minimum of one unique peptide per protein group. Carbamidomethylation of Cysteine was set as a fixed modification and oxidation of Methionine as well as N-terminal formylation and acetylation were set as variable modification. Percolator was used to estimate the number of false positive identifications and the results were filtered for a strict target false discovery rate (FDR) < 0.01.

The mass spectrometry proteomics data have been deposited to the ProteomeXchange Consortium (http://proteomecentral.proteomexchange.org) via the PRIDE partner repository^42^ with the dataset identifier PXD061441.

### Cryo-electron microscopy

For cryo-EM sample preparation, 4.5µl of the purified complex were applied to glow discharged Quantifoil 2/1 grids, blotted for 3.5s with force 4 in a Vitrobot Mark III (Thermo Fisher) at 100% humidity and 4°C, and plunge frozen in liquid ethane, cooled by liquid nitrogen. Cryo-EM data was acquired with an FEI Titan Krios transmission electron microscope using SerialEM software^43^. Movie frames were recorded at a nominal magnification of 29,000X using a K3 direct electron detector (Gatan). The total electron dose of ∼55 electrons per Å^2^ was distributed over 30 frames at a calibrated physical pixel size of 0.84 Å. Micrographs were recorded in a defocus range of −0.5 to −3.0 μm.

### Image processing, classification, and refinement

Cryo-EM micrographs of PSII isolated by TS-Psb27 were processed on the fly using the Focus software package^44^ if they passed the selection criteria (iciness < 1.05, drift 0.4 Å < x < 70 Å, defocus 0.5 µm < x < 5.5 µm, estimated CTF resolution < 6 Å). Micrograph frames were aligned using MotionCor2^45^ and the contrast transfer function (CTF) for aligned frames was determined using GCTF^46^. From 20,182 acquired micrographs 5,055,941 particles were picked using the Phosaurus neural network architecture from crYOLO^47^. Particles were extracted with a pixel box size of 256 scaled down to 64 using RELION 3.1^48^ and underwent several rounds of reference-free 2D classification At this stage, obvious non-particles (carbon edges, contamination, overlapping particles) and views that failed to align coherently were removed, as these species do not contribute meaningful signal and would otherwise reduce the resolution of downstream reconstructions. The remaining 3,100,431 selected particles were re-extracted with a box size of 320 and imported into Cryosparc 2.3^49^, given its GPU-accelerated algorithms enable substantially faster exploration of structural heterogeneity, an advantage when handling millions of particles and multiple potential intermediate states. After *ab initio* model generation and heterogeneous refinement particles of the Psb32-PSII like class were re-imported to RELION and further classified in 2D. Poorly aligned particles that did not contribute to meaningful reconstruction were discarded. Selected particles were down-sampled to a box size of 128 and refined against a consensus model. Using 3D focused classification without alignment gave rise to distinct classes with (199,653 particles) and without Psb32 (325,601), while discarding residual heterogeneous particles that did not converge to defined densities. Several rounds of refinement, CTF-refinement (estimation of anisotropic magnification, fit of per-micrograph defocus and astigmatism and beam tilt estimation) and Bayesian polishing^50^ yielded final models with an estimated resolution of ∼2.9 Å and ∼2.8 Å for PSII with Psb32 and without Psb32, respectively (Gold standard FSC analysis of two independent half-sets at the 0.143 cutoff). Local-resolution and 3D-FSC plots (Figure S3) were calculated using RELION and the “Remote 3DFSC Processing Server” web interface^51^, respectively.

### Atomic model construction

The 3.6 Å resolution X-ray crystal structure of monomeric PSII from *T. vestitus* with PDB-ID 3KZI^52^ was used as initial structural model that was docked as rigid body using Chimera^53^ into the obtained cryo-EM density maps of Psb32-PSII and Psb27-PSII. For all PSII complexes, the subunits and cofactors that had no corresponding density were removed. By highlighting the still unoccupied parts of the density maps, we identified the density regions that belong to subunits of the PSII that are not present in the initial structure and therefore need to be added to the model. For Psb32-PSII we assigned the unoccupied densities to the mass spectrometry identified subunits Psb27 and Psb32. For Psb27-PSII we assigned Psb27. For Psb32-PSII we deleted PsbO and PsbU from the initial structural model and for Psb27-PSII we deleted PsbO, PsbU, and PsbV. The 1.6 Å resolution X-ray crystal structure of isolated Psb27 from *T. vestitus* with PDB-ID 2Y6X^54^ was docked as rigid body into the unoccupied density assigned to Psb27 in all three complexes.

We first determined the atomic structure of isolated mature Psb32 (UniProt ID Q8DLS4) with a fused sfGFP at the N-terminus using X-ray crystallography based on data obtained earlier^55^. The initial model after determination of phases by BALBES^56^ was further refined using Phenix^57^ and Coot^58^ against the native dataset with a resolution of 2.12 Å. Refinement included individual B-factors and TLS refinement using TLS groups automatically defined by Phenix. Residues 0 to150 of the deposited coordinates (PDB-ID 8C7I), which correspond to the residues 30 to 180 in the Uniprot entry Q8DLS4, were then used for docking into the previously assigned cryo-EM density. Interestingly, there was still unoccupied density left reflecting the truncated C-terminal tail of Psb32 that could not be resolved by X-ray crystallography due to its high flexibility in the isolated form.

We predicted models of the complete Psb32 sequence (UniProt ID Q8DLS4) using AlphaFold2^59^ and AlphaFold3^60^. We also predicted the secondary structure through the meta server Bioinformatics Toolkit^61^ and CCTOP^62^. All results provide an almost identical length of the C-terminal α-helix. Therefore, we merged the C-terminus from residue 181 to 228 of the AlphaFold3 predicted model with the docked Psb32 X-ray crystal structure. Then, we performed interactive molecular dynamics flexible fitting (MDFF) to fit the C-terminus into the identified unoccupied density.

### Model Refinement

Our employed model refinement strategy is well established and was applied to construct structural models of several different complex molecular machines by employing different bioinformatics protein modelling and simulation methods to interpret cryo-EM. The initial model of the complexes described above were refined in real space against the respective cryo-EM density, and structural clashes were removed using MDFF^63^. MDFF simulations were prepared in VMD 1.9.4a35^64^ using QwikMD^65^ and the MDFF plugin. The simulations were carried out with NAMD 2.13^66^ employing the CHARMM36 force field. Secondary structure, cis peptide and chirality restraints were employed during 800 steps of minimization followed by a 40 ps MDFF simulation at 300K. Due to the employed restraints, only conformational changes of side chains and subunit movements compared to the initial structure are identified during the initial MDFF run. We checked the fit to density of the structure by calculating cross-correlation values of the backbone atoms.

After the initial MDFF runs, the cross-correlation check did not reveal any regions with significant deviation between model and density. Therefore, no further refinement of secondary structure elements was necessary. This fast convergence reflects that there are no large conformational differences between the model and the initial X-ray crystal structures, except for the presence of the further subunits. Then, all three models were used to initiate one final round of real-space refinement in Phenix^57^. Last, a final inspection of steric clashes and chemical correctness e.g. of side chain head group orientations is performed using the prepare and inspect algorithms implemented in MAXIMOBY (CHEOPS, Germany).

### Structure Analysis

The atomic interaction pattern between the subunits were identified and analyzed using the MAXI-MOBY (CHEOPS, Germany) contact matrix algorithm and the VMD plugin PyContact^67^. Structural difference between structures were identified and highlighted through calculation of the residue wise root mean square deviation of the backbone Cα atoms.

### Phylogenetic Analysis

For the phylogenetic analysis, Psb32 and Psb32-like sequences from Streptophyta, Chlorophyta, Cyanobacteria and Bacteria were collected from the NCBI database (June 2024) using BLAST. Sequences were aligned with MAFFT 7.110 using the iterative refinement method with global pairwise alignment information (G-INS-i) and an offset value (gap extension penalty) of 2. Cyanobacteria encode a second psb32-like paralog with a long insertion near the C-terminus. These sequences were aligned separately and then combined with the other sequences in a profile alignment using the merge function of MAFFT. The resulting alignment was manually curated by removing lineage-specific insertions.

A maximum likelihood phylogeny was inferred using IQ-TREE 2.0.3. Using ModelFinder restricted to nuclear amino acid models the best fit was determined to be the general matrix WAG with empirical amino acid frequencies, a proportion of invariable sites, and the discrete gamma model with 4 rate categories to account for rate heterogeneity across sites (WAG+F+I+G4). The resulting tree was rooted with sequences from Margulisbacteria which form a distinct and monophyletic outgroup. Standard non-parametric (Felsenstein) bootstraps were calculated in 100 replicates using IQ-TREE 2.0.3 to assess the robustness of the phylogenetic inference.

## Supplementary information

### Supplementary discussion

#### PSII Preparation and biochemical characterization

A *Thermosynechococcus vestitus* BP-1 mutant line was created, in which the native Psb27-sequence was extended by an N-terminal Twin-Strep tag after the lipid-modified cysteine (supplementary Fig. S1). Psb27-containing PSII complexes were purified by Strep-tag affinity chromatography (ST-AC) after thylakoid-extraction with 1.2% (w/v) n-dodecyl-ß-d-maltoside (DDM) and 0.5% (w/v) Na-cholate (supplementary Fig. S2A). Most of the PSII reflects monomeric PSII with slightly different sizes, which indicates a mixture of different Psb27-containing PSII (supplementary Fig. S2B). Additionally, low amounts of dimeric PSII and PSI trimers are present. The separation into the subunits by SDS-PAGE revealed stoichiometric amounts of all major subunits and TS-Psb27 (supplementary Fig. S2C). Moreover, the low-molecular-weight proteins (LMWPs) PsbE, PsbF, PsbH, PsbI, PsbK, PsbL, PsbM, PsbT, PsbX, PsbZ, Psb30 and traces of PsbY were detected by MALDI-ToF-MS (supplementary Tab. S1). Most noteworthy, the PsbJ subunit is present, which is absent in the recent cryo-EM structures of Psb27-containing PSII complexes^1,2^. Psb34 and Psb28 were also detected by MS, although Psb28 was not clearly visible SDS-PAGE due to the intensive band of the TS-Psb27 (Supplementary Tab. S1, supplementary Data 1). The unknown band at around 25 kDa was assigned to Psb32 (Tll0404, Q8DLS4) by single-band LC-MS/MS (Supplementary Data 2). Similar to observations by Liu et al.^3^ which used a His-tagged Psb27, PsbO was co-purified in sub-stoichiometric amounts. Several other proteins were identified by MS, including CyanoP, CyanoQ, PsbV, PsbU, FtsH as well as subunits from the membrane complexes NDH-1 and PSI (Supplementary Data 1). The Psb27-containing PSII fraction shows only residual oxygen evolution (363 ± 29 µmol O_2_·(mg Chl·h)^-1^) compared to active PSII (5,000-6,000 µmol O_2_·(mg Chl·h)^-1^) from *T. vestitus* obtained by similar preparations^4–6^.

### Supplementary tables and figures

**Table S1:**
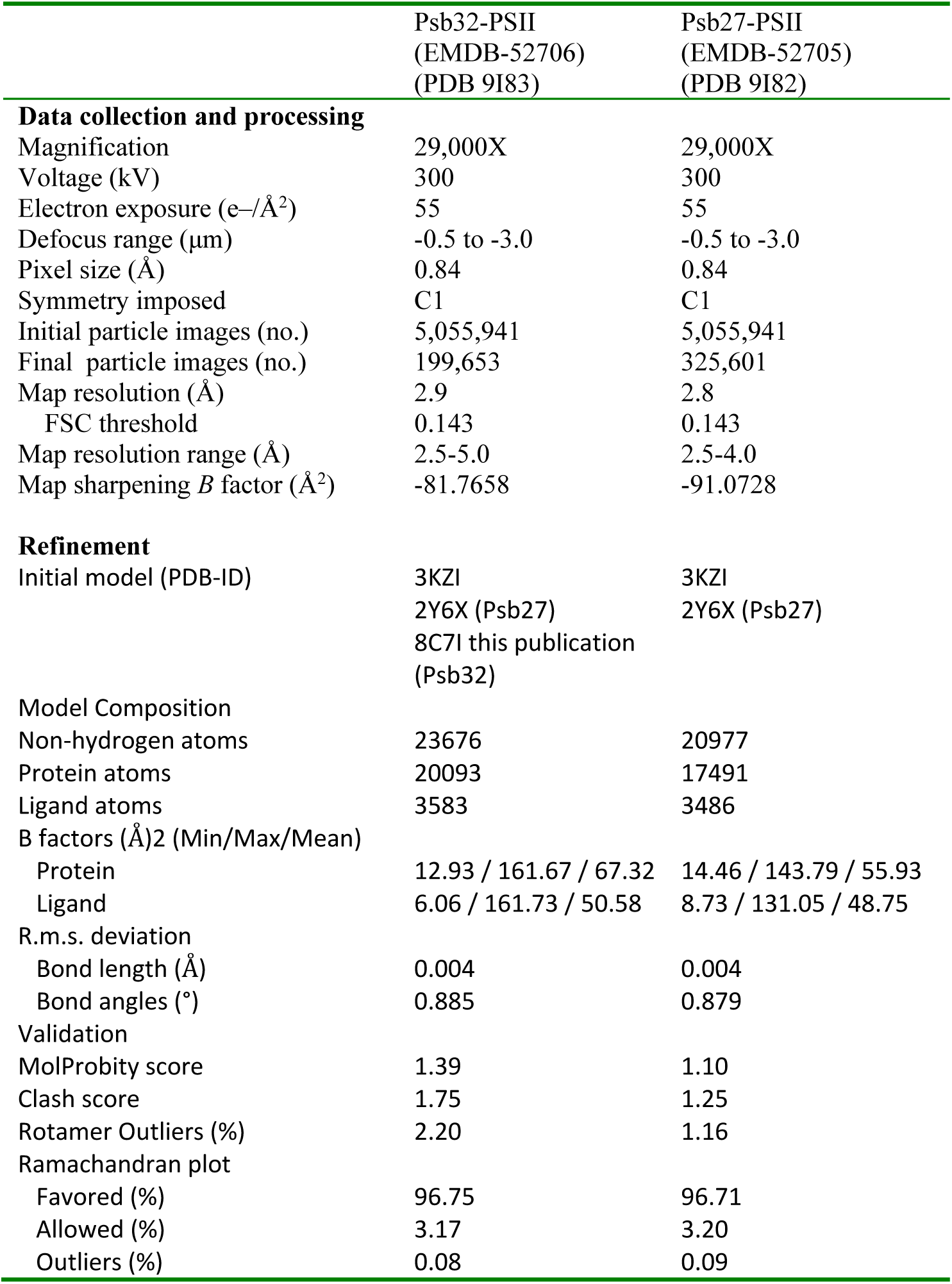
Cryo-EM data collection, refinement, and validation statistics.

|  | Psb32-PSII<br>(EMDB-52706)<br>(PDB 9I83) | Psb27-PSII<br>(EMDB-52705)<br>(PDB 9I82) |
| --- | --- | --- |
| **Data collection and processing** |  |  |
| Magnification | 29,000X | 29,000X |
| Voltage (kV) | 300 | 300 |
| Electron exposure (e-/Å<sup>2</sup>) | 55 | 55 |
| Defocus range (µm) | -0.5 to -3.0 | -0.5 to -3.0 |
| Pixel size (Å) | 0.84 | 0.84 |
| Symmetry imposed | C1 | C1 |
| Initial particle images (no.) | 5,055,941 | 5,055,941 |
| Final particle images (no.) | 199,653 | 325,601 |
| Map resolution (Å) | 2.9 | 2.8 |
| FSC threshold | 0.143 | 0.143 |
| Map resolution range (Å) | 2.5-5.0 | 2.5-4.0 |
| Map sharpening *B* factor (Å<sup>2</sup>) | -81.7658 | -91.0728 |
| **Refinement** |  |  |
| Initial model (PDB-ID) | 3KZI<br>2Y6X (Psb27)<br>8C7I this publication<br>(Psb32) | 3KZI<br>2Y6X (Psb27) |
| **Model Composition** |  |  |
| Non-hydrogen atoms | 23676 | 20977 |
| Protein atoms | 20093 | 17491 |
| Ligand atoms | 3583 | 3486 |
| B factors (Å)<sup>2</sup> (Min/Max/Mean) |  |  |
| Protein | 12.93 / 161.67 / 67.32 | 14.46 / 143.79 / 55.93 |
| Ligand | 6.06 / 161.73 / 50.58 | 8.73 / 131.05 / 48.75 |
| **R.m.s. deviation** |  |  |
| Bond length (Å) | 0.004 | 0.004 |
| Bond angles (°) | 0.885 | 0.879 |
| **Validation** |  |  |
| MolProbity score | 1.39 | 1.10 |
| Clash score | 1.75 | 1.25 |
| Rotamer Outliers (%) | 2.20 | 1.16 |
| **Ramachandran plot** |  |  |
| Favored (%) | 96.75 | 96.71 |
| Allowed (%) | 3.17 | 3.20 |
| Outliers (%) | 0.08 | 0.09 |

**Table S2:** Crystal structure statistics. Values in parentheses are for the highest resolution shell.

|  |  |
| --- | --- |
|  | sgpsb32_6 |
| Diffraction Source | SLS PXII |
| Wavelength (Å) | 0.975107 |
| Temperature (K) | 100 |
| Detector | PILATUS 6M |
| Crystal-to-detector distance (mm) | 435 |
| Rotation range per image (°) | 0.25 |
| Total rotation range (°) | 325 |
| Exposure time per image (s) | 0.1 |
| Space group | *P*6<sub>1</sub>22 |
| a, b, c (Å) | 145.65, 145.65, 91.81 |
| α, β, γ (°) | 90, 90, 120 |
| Resolution range (Å) | 45.68-2.12 (2.18-2.12) |
| Total No. of reflections | 1158546 (88765) |
| No. of unique reflections | 32795 (2370) |
| Completeness (%) | 99.9 (100) |
| Multiplicity | 35.3 (37.5) |
| ⟨I/σ(I)⟩ | 24.87 (1.01) |
| R<sub>meas</sub> (%) | 13.0 (675.2) |
| CC1/2 | 100 (56.3) |
| Overall B factor from Wilson plot (Å<sup>2</sup>) | 64.1 |
| Refinement statistics |  |
| Reflections used in refinement | 32779 |
| Reflections used for R-free | 1639 |
| R-work | 0.2504 |
| R-free | 0.2787 |
| CC(work) | 0.944 |
| CC(free) | 0.948 |
| Number of non-hydrogen atoms | 3165 |
| macromolecules | 3109 |
| ligands | 39 |
| solvent | 33 |
| Protein residues | 398 |
| RMS(bonds) | 0.004 |
| RMS(angles) | 0.58 |
| Ramachandran favored (%) | 96.43 |
| Ramachandran allowed (%) | 2.81 |
| Ramachandran outliers (%) | 0.77 |
| Rotamer outliers (%) | 1.17 |
| Clashscore | 5.18 |
| Average B-factor | 86.89 |
| macromolecules | 87.21 |
| ligands | 70.95 |
| solvent | 68.27 |
| Number of TLS groups | 2 |

**Table S3.** Coordinating interactions of Psb32 with the other PSII subunits. Listed are all specific contacts between Psb32 and the rest of the complex as well as the number of van der Waals contacts. Coloring according to figure 1.

| Subunit 1 | Subunit 2 | Residue 1 | Residue 2 | Contact Type |
| --- | --- | --- | --- | --- |
| Psb32 | Psb27 | Arg82 | Ser102 | H-Bond |
| Psb32 | Psb27 | Arg82 | Ser103 | H-Bond |
| Psb32 | Psb27 | Arg82 | Pro105 | Specific<br>Van der Waals |
| Psb32 | Psb27 | Asp84 | Asn106 | H-Bond |
| Psb32 | Psb27 | Asp84 | Arg107 | H-Bond |
| Psb32 | Psb27 | Tyr85 | Tyr104 | Specific<br>Van der Waals |
| Psb32 | Psb27 | 4x |  | Van der Waals |
| Psb32 | PsbV | Val118 | Arg131 | H-Bond |
| Psb32 | PsbV | Arg152 | Asp137 | H-Bond |
| Psb32 | PsbV | Glu184 | Lys50 | H-Bond |
| Psb32 | PsbV | Glu189 | Glu28 | H-Bond |
| Psb32 | PsbV | 3x |  | Van der Waals |
| Psb32 | PsbE | Tyr191 | Pro52 | Specific<br>Van der Waals |
| Psb32 | PsbE | Tyr191 | Asp53 | H-Bond |
| Psb32 | PsbE | Lys192 | Arg51 | H-Bond |
| Psb32 | PsbE | Lys194 | Asp45 | H-Bond |
| Psb32 | PsbE | Thr197 | Thr40 | H-Bond |
| Psb32 | PsbE | Tyr224 | Arg18 | Specific<br>Van der Waals |
| Psb32 | PsbE | 25x |  | Van der Waals |
| Psb32 | PsbF | Thr190 | Phe42 | H-Bond |
| Psb32 | PsbF | Thr190 | Gln44 | H-Bond |
| Psb32 | PsbF | Thr190 | Gln44 | H-Bond |
| Psb32 | PsbF | Tyr221 | Phe16 | Specific<br>Van der Waals |
| Psb32 | PsbF | 3x |  | Van der Waals |

**Table S4.** Coordinating interactions of the D2 C-terminal tail in Psb32-PSII and the mature PSII complex. Listed are all contacts between the last five C-terminal residues of D2 and their interacting subunits within the immature Psb32-PSII assembly intermediate (PDB-ID: 9I83) and mature PSII (PDB-ID: 4PJ0). Coloring according to figure 1.

| Subunit 1 | Subunit 2 | Residue 1<br>(in-file Numbering) | Residue 2<br>(in-file Numbering) | Contact Type |
| --- | --- | --- | --- | --- |
| **Psb32-PSII** |  |  |  |  |
| D2 | D1 | Pro347 | Met331 | Van der Waals |
| D2 | CP47 | Pro347 | Phe383 | Specific<br>Van der Waals |
| D2 | D1 | Arg348 | Val330 | Van der Waals |
| D2 | D1 | Gly349 | Val330 | H-Bond |
| D2 | D1 | Gly349 | Arg334 | Van der Waals |
| D2 | CP47 | Asn350 | Arg384 | H-Bond |
| D2 | CP47 | Ala351 | Phe383 | Van der Waals |
| D2 | CP47 | Ala351 | Arg384 | Van der Waals |
| D2 | CP47 | Ala351 | Arg384 | Van der Waals |
| D2 | CP47 | Leu352 | Phe383 | Specific<br>Van der Waals |
| D2 | CP47 | Leu352 | Arg383 | Van der Waals |
| **Mature PSII** |  |  |  |  |
| D2 | CP47 | Pro347 | Phe383 | Van der Waals |
| D2 | PsbU | Arg348 | Lys104 | H-Bond |
| D2 | D1 | Gly349 | Val330 | H-Bond |
| D2 | D1 | Gly349 | Arg334 | Van der Waals |
| D2 | CP47 | Gly349 | Arg384 | H-Bond |
| D2 | D1 | Asn350 | Arg334 | H-Bond |
| D2 | D1 | Asn350 | His337 | H-Bond |
| D2 | PsbO | Asn350 | Asn155 | H-Bond |
| D2 | D1 | Ala351 | His337 | Van der Waals |
| D2 | D1 | Ala351 | Asn338 | Van der Waals |
| D2 | PsbU | Ala351 | Gly101 | H-Bond |
| D2 | PsbU | Ala351 | Tyr103 | Van der Waals |
| D2 | PsbU | Ala351 | Tyr103 | Van der Waals |
| D2 | D1 | Leu352 | His332 | Van der Waals |
| D2 | D1 | Leu352 | His337 | Specific<br>Van der Waals |
| D2 | D1 | Leu352 | His337 | Van der Waals |

**Table S5:** Proteins identified by MALDI-ToF and LC-MS/MS.

| **Accession** | **Gene name** | **Description** | **Detected by LC-MS** | **Detected by MALDI** |
| --- | --- | --- | --- | --- |
| P0A444 | PsbA1 | Photosystem II protein D1 1 | + |  |
| P0A446 | PsbA2 | Photosystem II protein D1 2 | + |  |
| Q8DIQ1 | PsbB | Photosystem II CP47 reaction center protein | + |  |
| Q8DIF8 | PsbC | Photosystem II CP43 reaction center protein | + |  |
| Q8CM25 | PsbD1 | Photosystem II D2 protein | + |  |
| Q8DIP0 | PsbE | Cytochrome *b*<sub>559</sub> subunit alpha | + | + |
| Q8DIN9 | PsbF | Cytochrome *b*<sub>559</sub> subunit beta | + | + |
| Q8DJ43 | PsbH | Photosystem II reaction center protein H | + | + |
| Q8DJZ6 | PsbI | Photosystem II reaction center protein I | + | + |
| P59087 | PsbJ | Photosystem II reaction center protein J | + | + |
| Q9F1K9 | PsbK | Photosystem II reaction center protein K | + | + |
| Q8DIN8 | PsbL | Photosystem II reaction center protein L | + | + |
| Q8DHA7 | PsbM | Photosystem II reaction center protein M | + | + |
| P0A431 | PsbO | Photosystem II manganese-stabilizing polypeptide | + |  |
| Q8DH84 | PsbP | PsbP protein | + |  |
| Q8DIQ0 | PsbT | Photosystem II reaction center protein T | + |  |
| P0A386 | PsbV | Cytochrome *c*<sub>550</sub> | + | + |
| Q9F1R6 | PsbX | Photosystem II reaction center X protein | + |  |
| Q8DKM3 | PsbY | Photosystem II protein Y | + | + |
| Q8DHJ2 | PsbZ | Photosystem II reaction center protein Z | + | + |
| **PSII associated** |  |  |  |  |
| Q8DG60 | TS-Psb27 | Photosystem II lipoprotein Psb27 | + |  |
| Q8DLJ8 | Psb28 | Photosystem II reaction center Psb28 protein | + |  |
| Q8DLS4 | tll0404 | Tll0404 protein (Psb32) | + |  |
| Q8DMP8 | tsl0063 | Tsl0063 protein (Psb34) | + | + |
| Q8DJE2 | PsbV2 | Cytochrome *c*<sub>550</sub>-like protein | + |  |
| Q8DJI1 | Psb30 | Photosystem II reaction center protein Ycf12/Psb30 |  | + |

**Figure S1:**
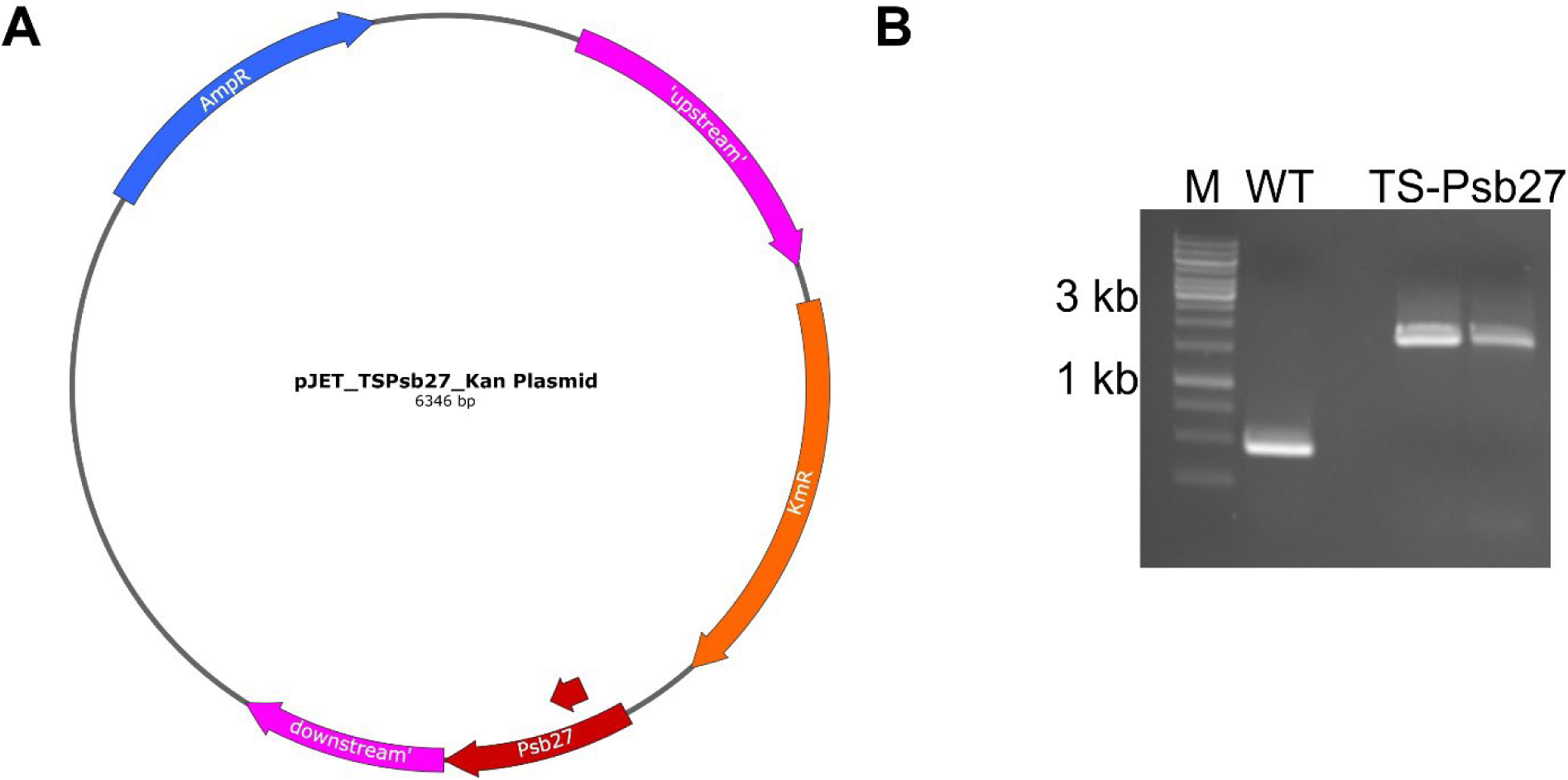
Construction of TwinStrep-tagged Psb27. **A)** Plasmid map used for the transformation. The plasmid contains resistance cassettes against ampicillin (blue) and kanamycin (orange). The up- and downstream regions are shown in pink and the Psb27 sequence and the TS-Tag (small arrow) are shown in red. **B)** Agarose gel electrophoresis of the PCR-Products of *T. vestitus* BP-1 (WT) and two TS-Psb27 mutants (see methods for primer details). A size standard (M), Thermo Scientific GeneRuler 1 kb DNA Ladder was applied.

**Figure S2:**
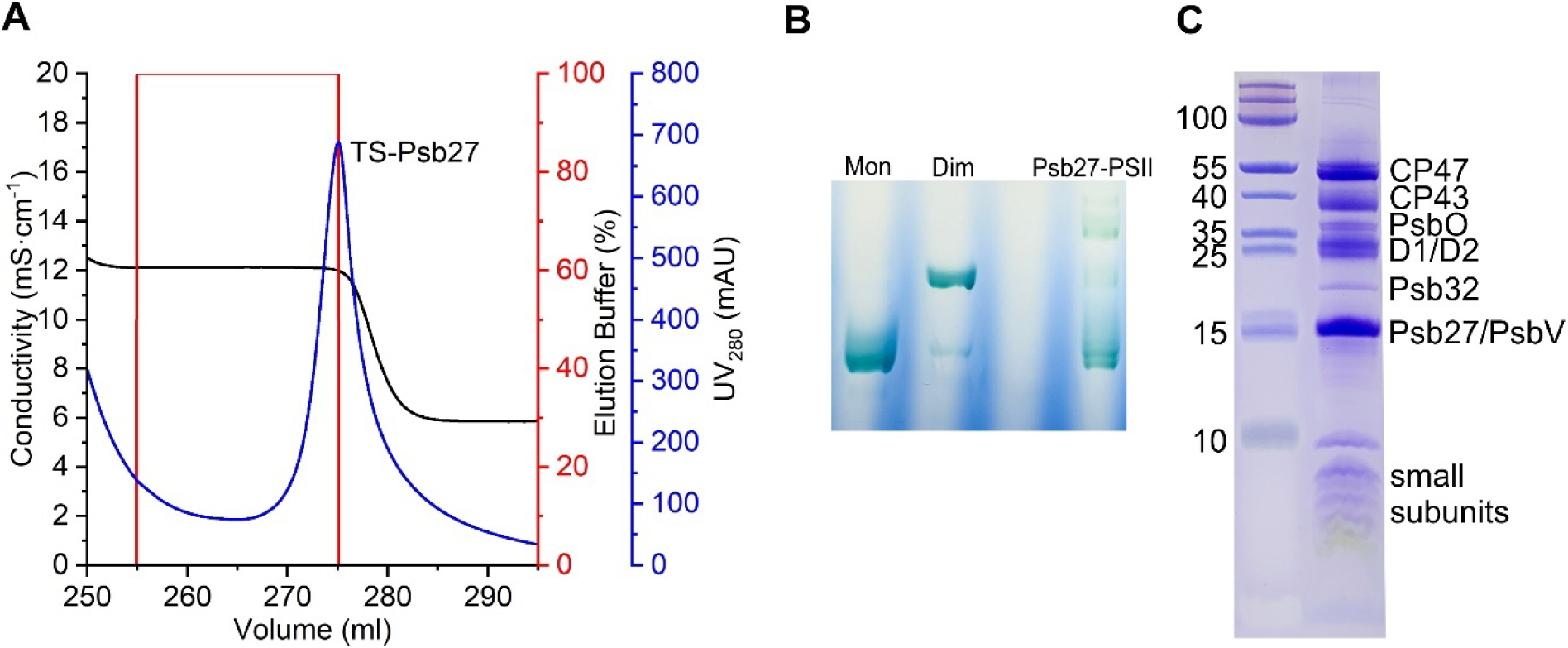
Preparation of PSII with TS-Psb27. **A)** Strep-Tag affinity chromatography (ST-AC) elution peak. The UV-absorption is shown in blue, the gradient of the elution buffer in red, and the conductivity in blue. The graphs shown are representative of several individual preparations. **B)** BN-PAGE and **C)** SDS-PAGE of purified PSII. A size standards (M), Thermofisher PageRuler #26616 was used for SDS-PAGE and PSII monomer/dimer purified by TS-tagged CP43 for BN-PAGE^5^.

**Figure S3:**
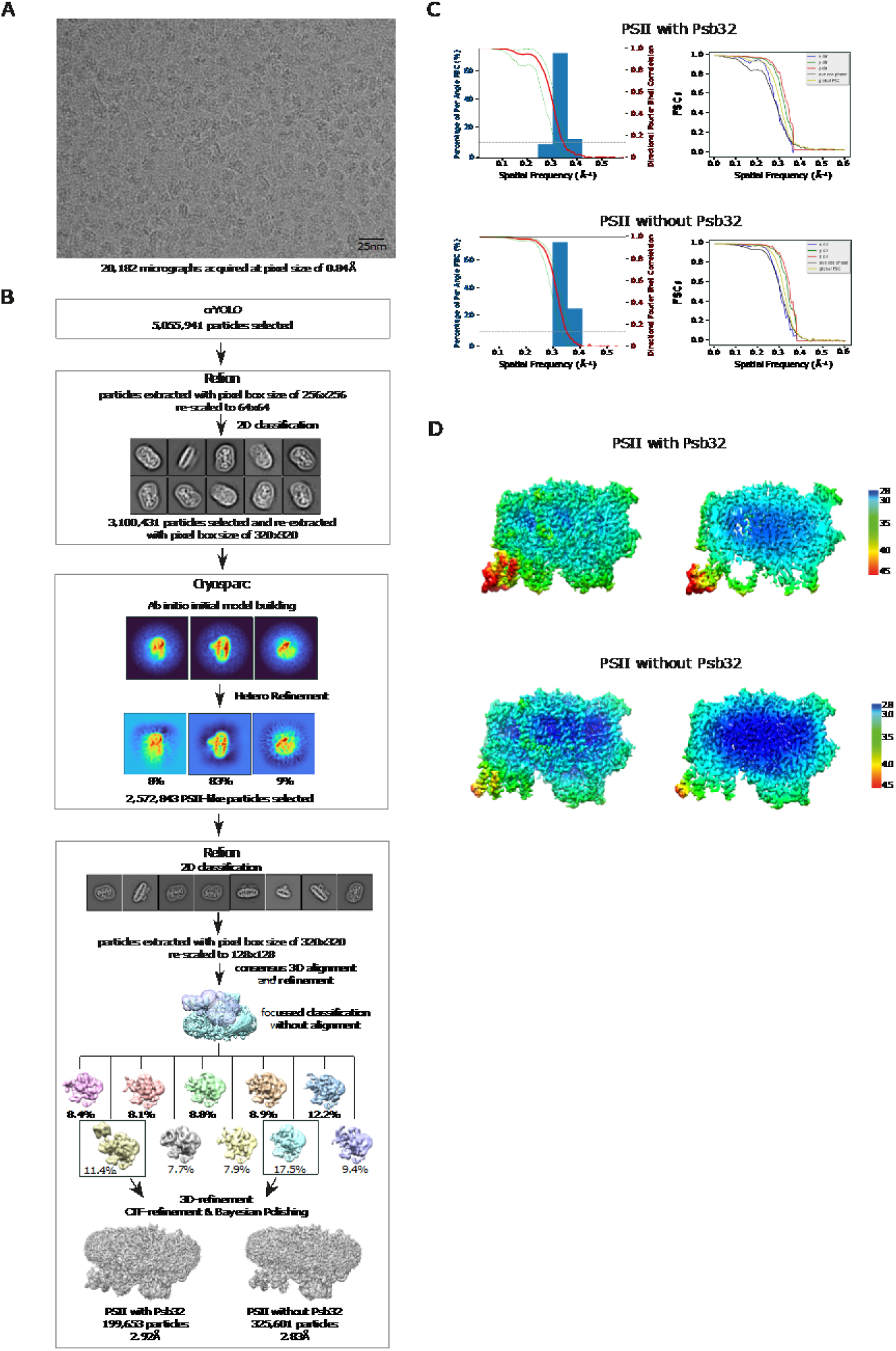
Cryo-EM data collection and processing scheme. **A)** Exemplary micrograph acquired on K3 (Gatan, 5760×4092 pixel) showing particle distribution on Quantifoil 2/1 grid-holes. **B)** Schematic of the image processing workflow. Particles were picked with crYOLO and extracted with RELION. After initial 2D classification, selected particles were used for ab initio model building and 3D-classification in Cryosparc. Particles belonging to the class resembling an intact PSII-particle were further classified in RELION (2D as well as focused classification using a mask). Classes resulting from focused refinement were refined and analyzed individually (not shown). Representatives of each identified unique conformer (with and without additional density in the mask region) were refined and particles belonging to the resulting two densities were subjected to CTF-refinement and Bayesian polishing before the final refinement. **C)** Plots showing the global (left plot) and directional (right plot) resolution for the two densities obtained in B), calculated using the “Remote 3DFSC Processing Server” web interface. For PSII with Psb32 and PSII without Psb32 a sphericity score of 0.947 and 0.970 and a global resolution (FSC-threshold = 0.143) of 2.99A and 2.86A were calculated, respectively. **D)** Local resolution as calculated by RELION mapped on the refined densities (left: surface view, right: cut-open view of central section).

**Figure S4:**
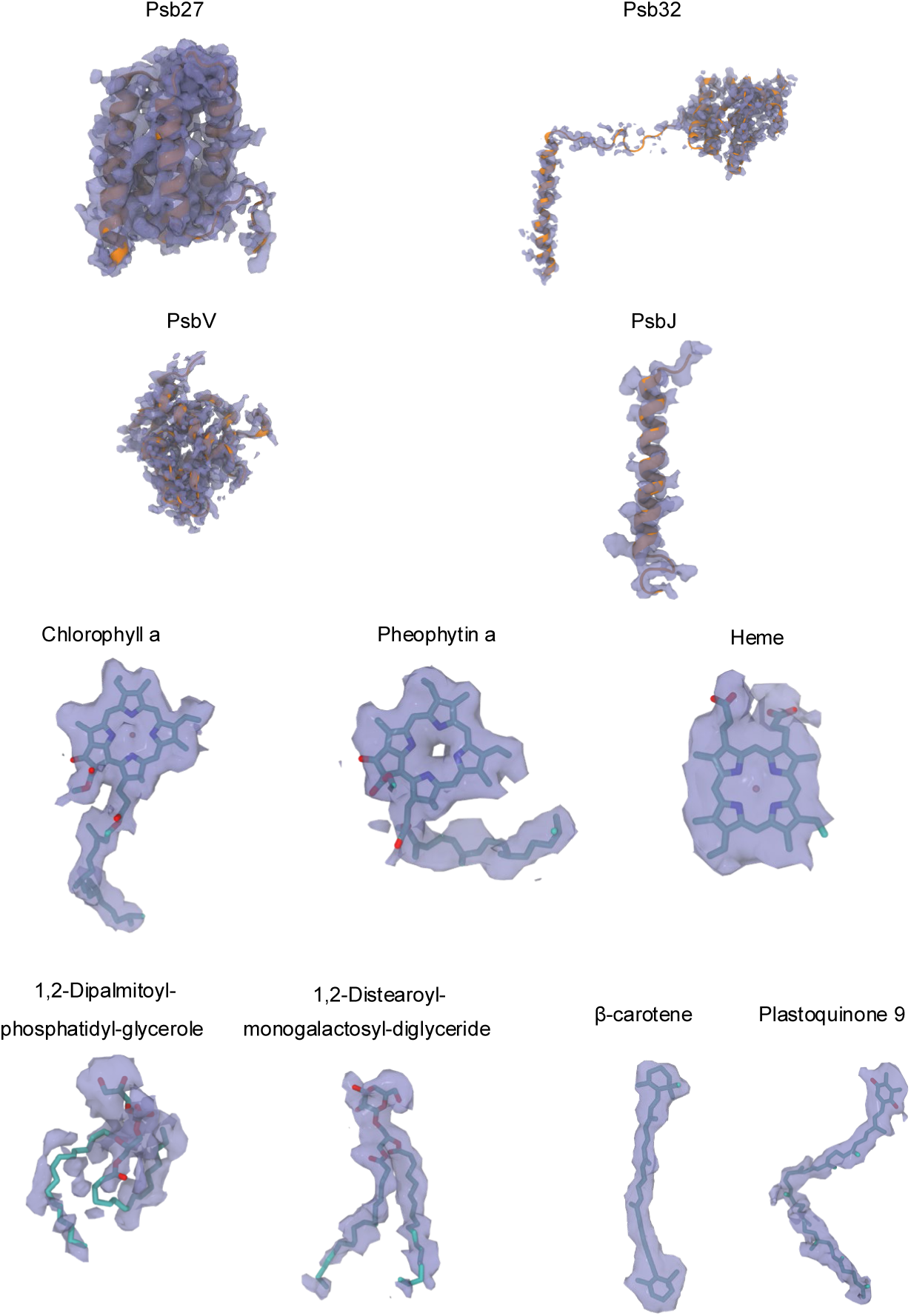
Comparison of structural model and density for Psb32-PSII. Fit of the structural model into the density map (transparent blue) for selected key subunits and cofactors of Psb32-PSII. The resolution is sufficient for accurate modeling of subunits and cofactors.

**Figure S5:**
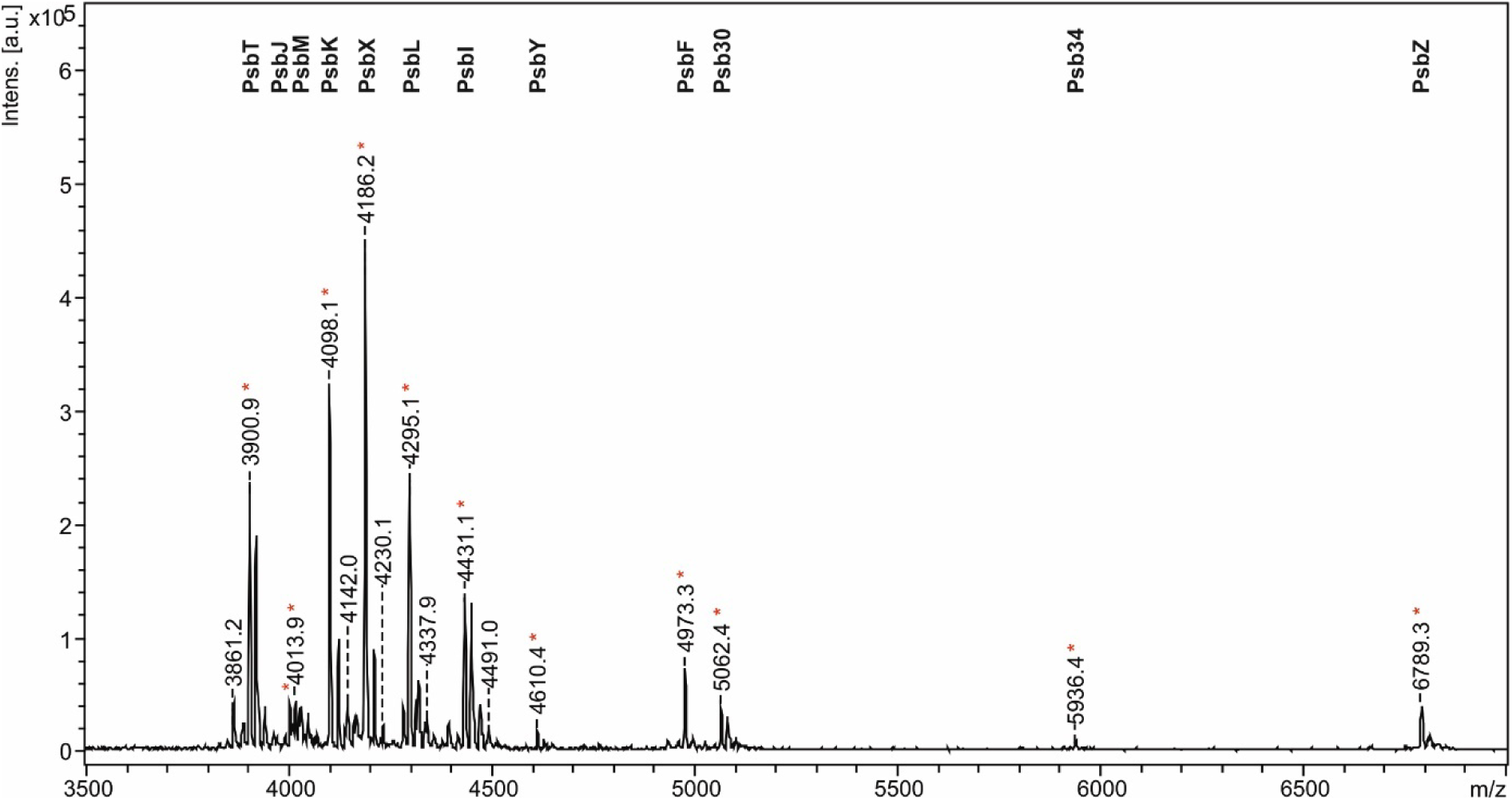
MALDI-ToF-MS spectra of the small proteins of PSII at the m/z range from 3,500 to 7,000. Annotation of the peaks was done according to^6–8^. Stars indicate modified peptides.

**Figure S6:**
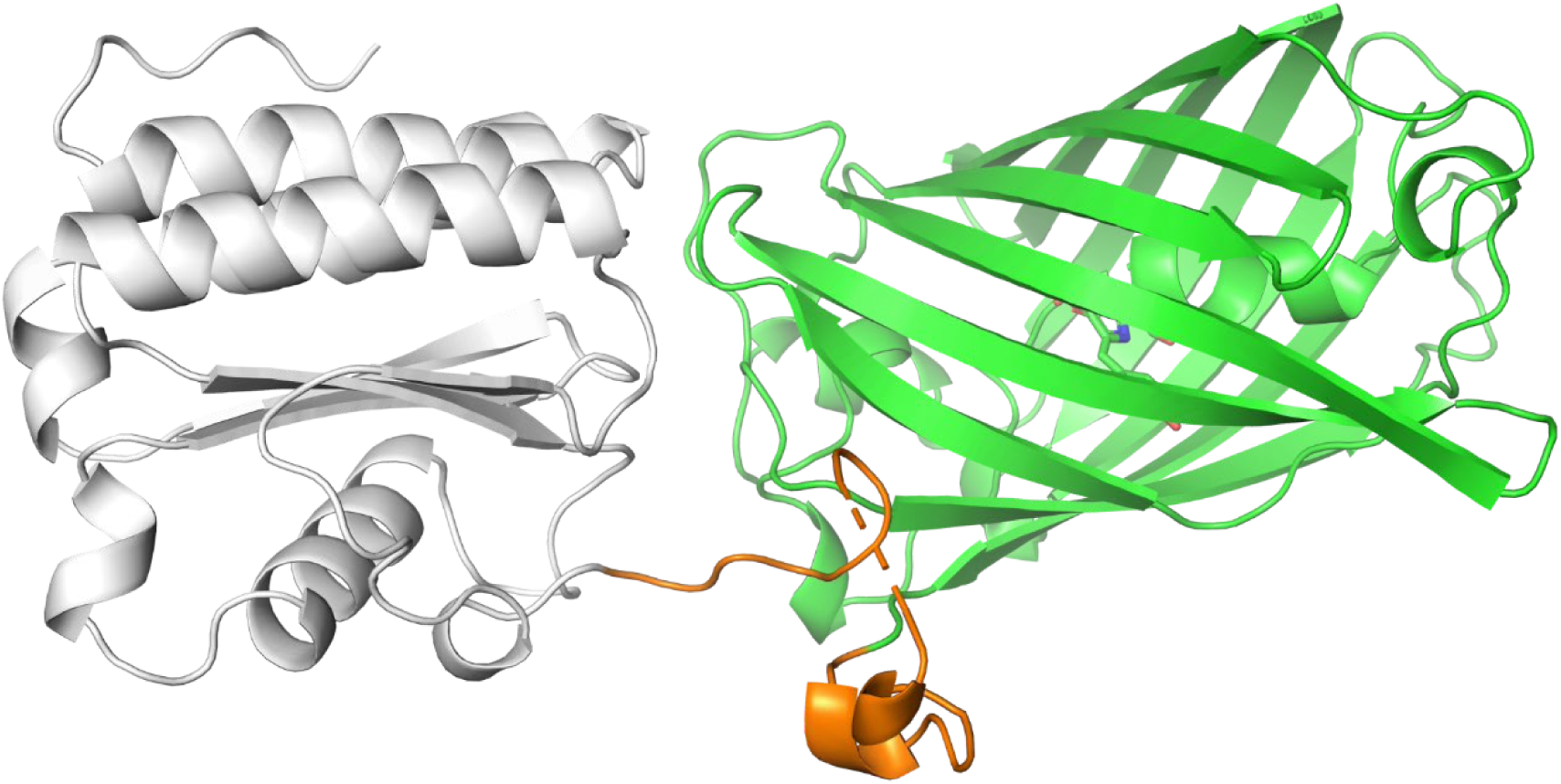
Psb32 crystal structure. Cartoon representation of the structure of the sfGFP-Psb32 fusion protein. The sfGFP is colored in green, Psb32 (residues 0 to 150 in PDB-ID 8C7I and residues 30-180 from Uniprot ID Q8DLS4) colored in grey. The partially disordered linker region is shown in orange.

**Figure S7:**
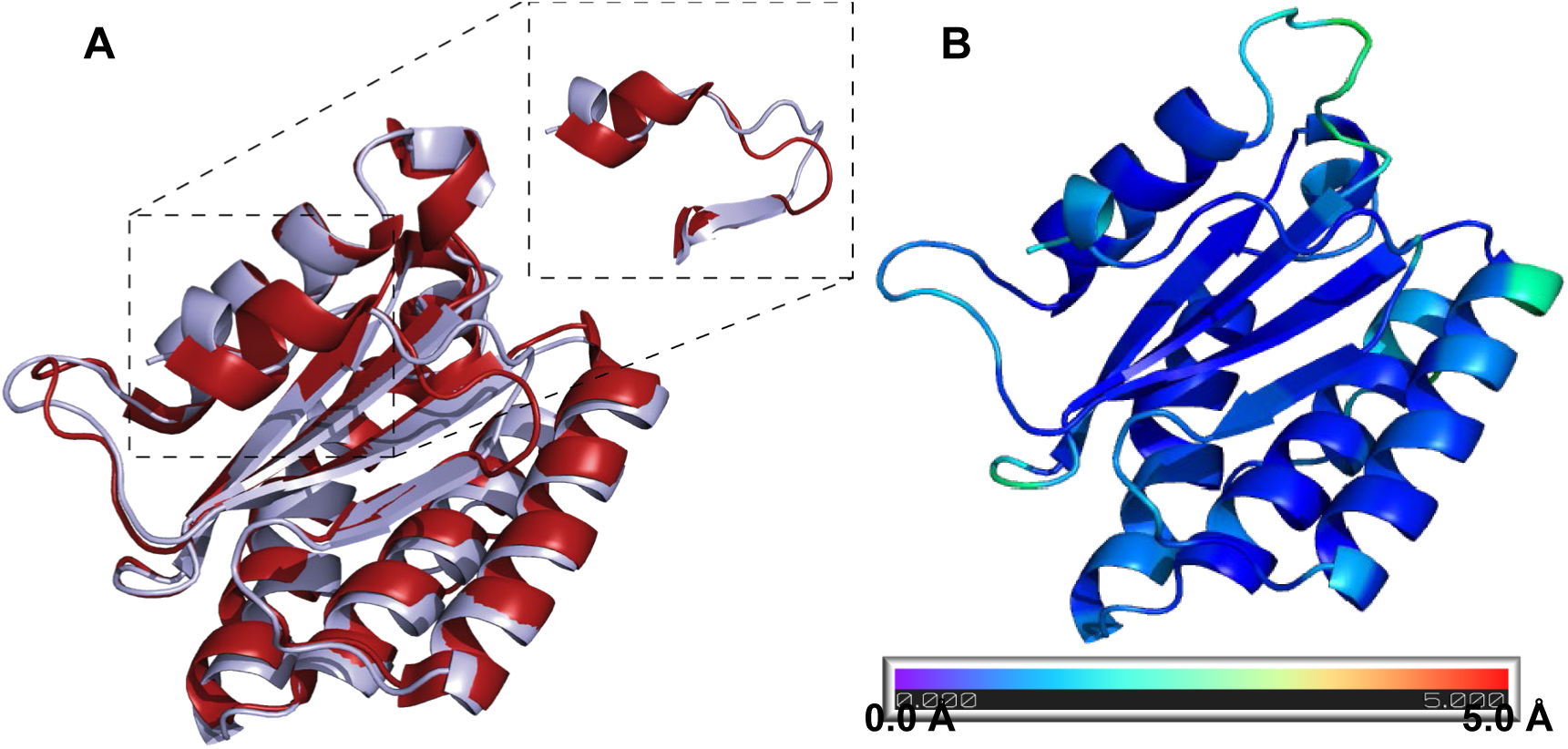
Comparison of Psb32 X-ray crystal structures. **A)** Cartoon representation of the here resolved X-ray crystal structure of Psb32 from *Thermosynechococcus vestitus* (light-blue, PDB-ID 8C7I) aligned to its *Arabidopsis thaliana* ortholog (red, PDB-ID 3PVH^9^). The structures are nearly identical, except for the extended C-terminus, which differs significantly in amino acid sequence. **B)** Depiction of the X-ray crystal structure of Psb32 colored according to the Cɑ root mean square deviation (RMSD) compared to the resolved globular domain within the cryo-EM model. The coloring ranges from 0 Å RMSD (deep blue) to 5 Å RMSD between these models.

**Figure S8:**
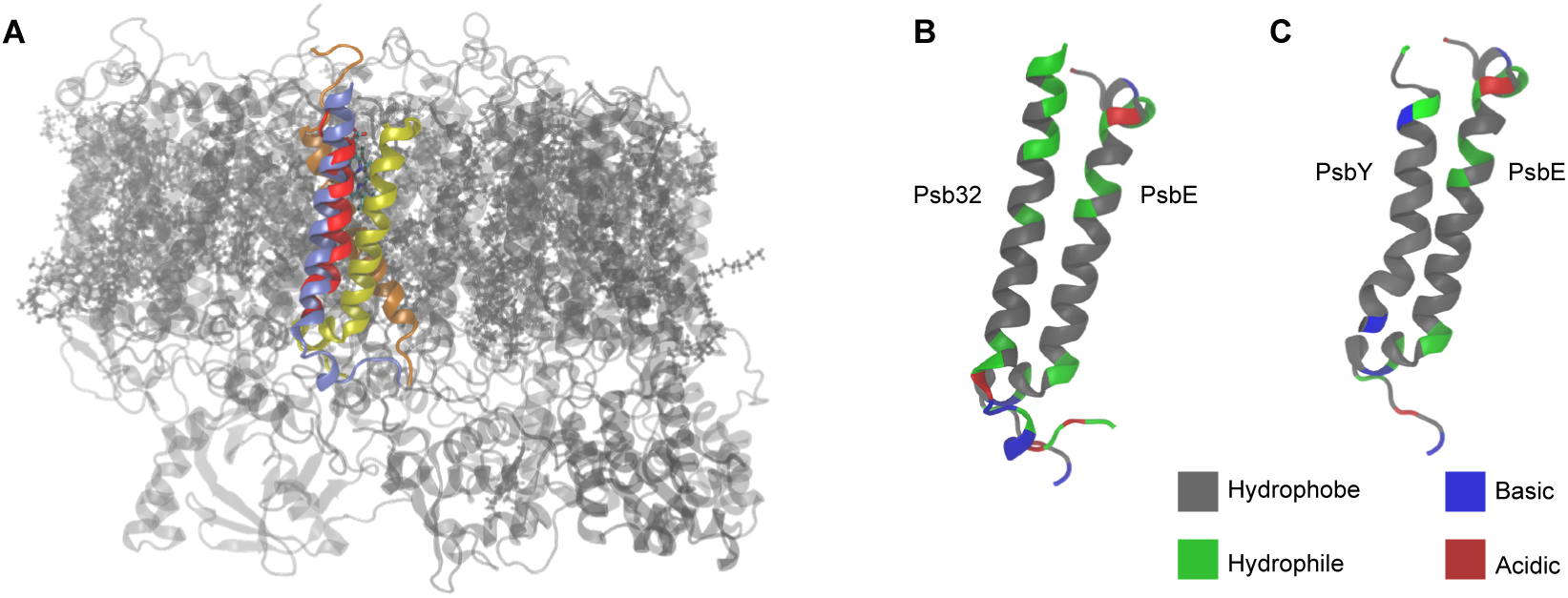
Assembly of Psb32 C-terminal Helix. **A)** Superposition of the mature PSII (PDB:4PJ0)^10^ with Psb32-PSII shows that the C-terminal helix of Psb32 (light blue) occupies a similar position as PsbY (red), both shielding the heme of Cyt *b*_559_. **B, C)** The different residue types of amino acids at the contact interface of are highlighted. **C)** Amino acid residue types at the contact interfaces of Psb32/PsbE and PsbY/PsbE, respectively. The abundance of hydrophobic residues and lack of compatible ionic or hydrophilic residues implies a mostly hydrophobic interaction for both Psb32 and PsbY.

**Figure S9:**
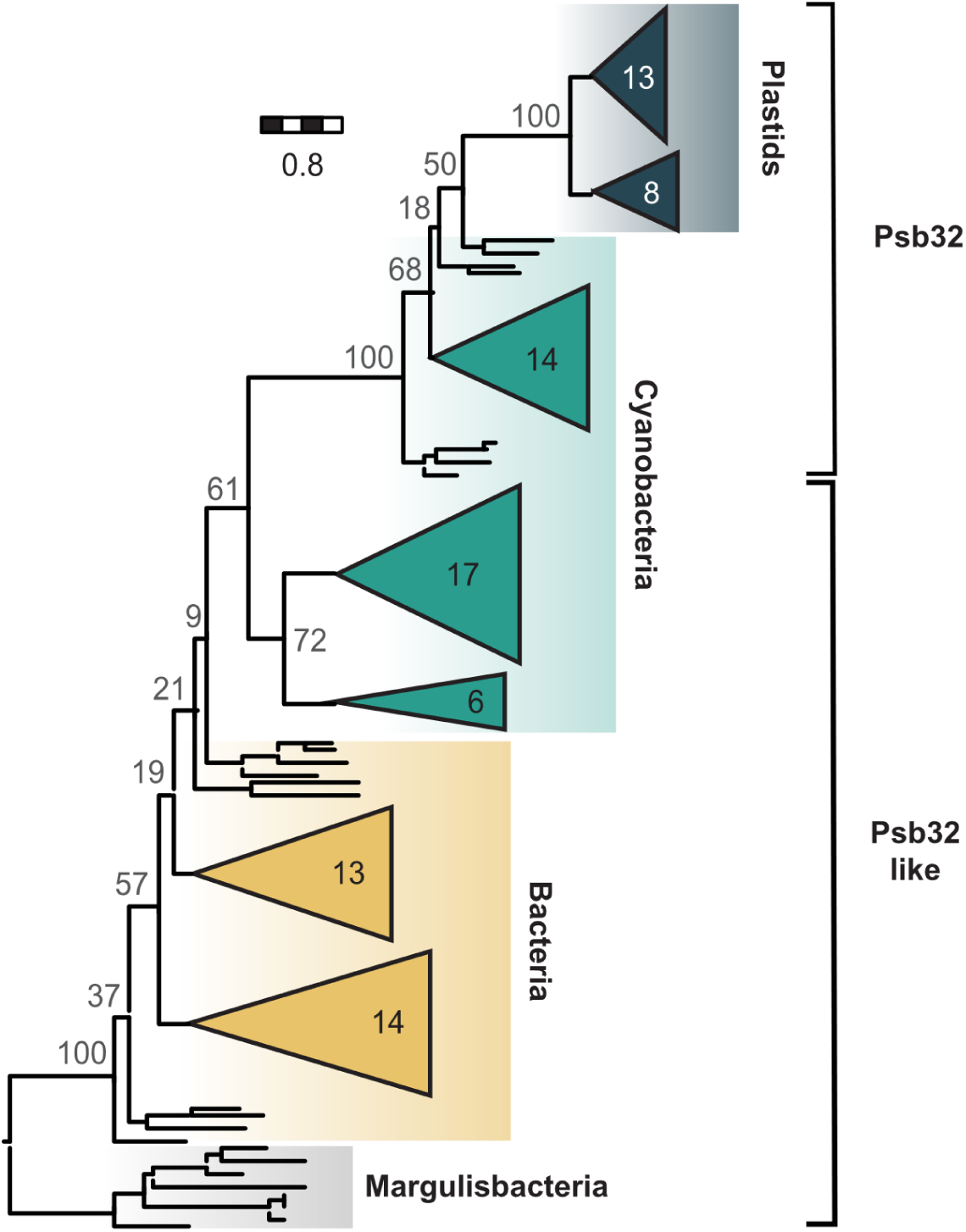
Phylogeny of Psb32 and Psb32-like homologues. Grey numbers next to nodes give non-parametric (Felsenstein) bootstraps, numbers in clades denote number of taxa per clade. The scale bar indicates how many amino acid substitutions happen on average per site along the horizontal branches.

**Figure S10:**
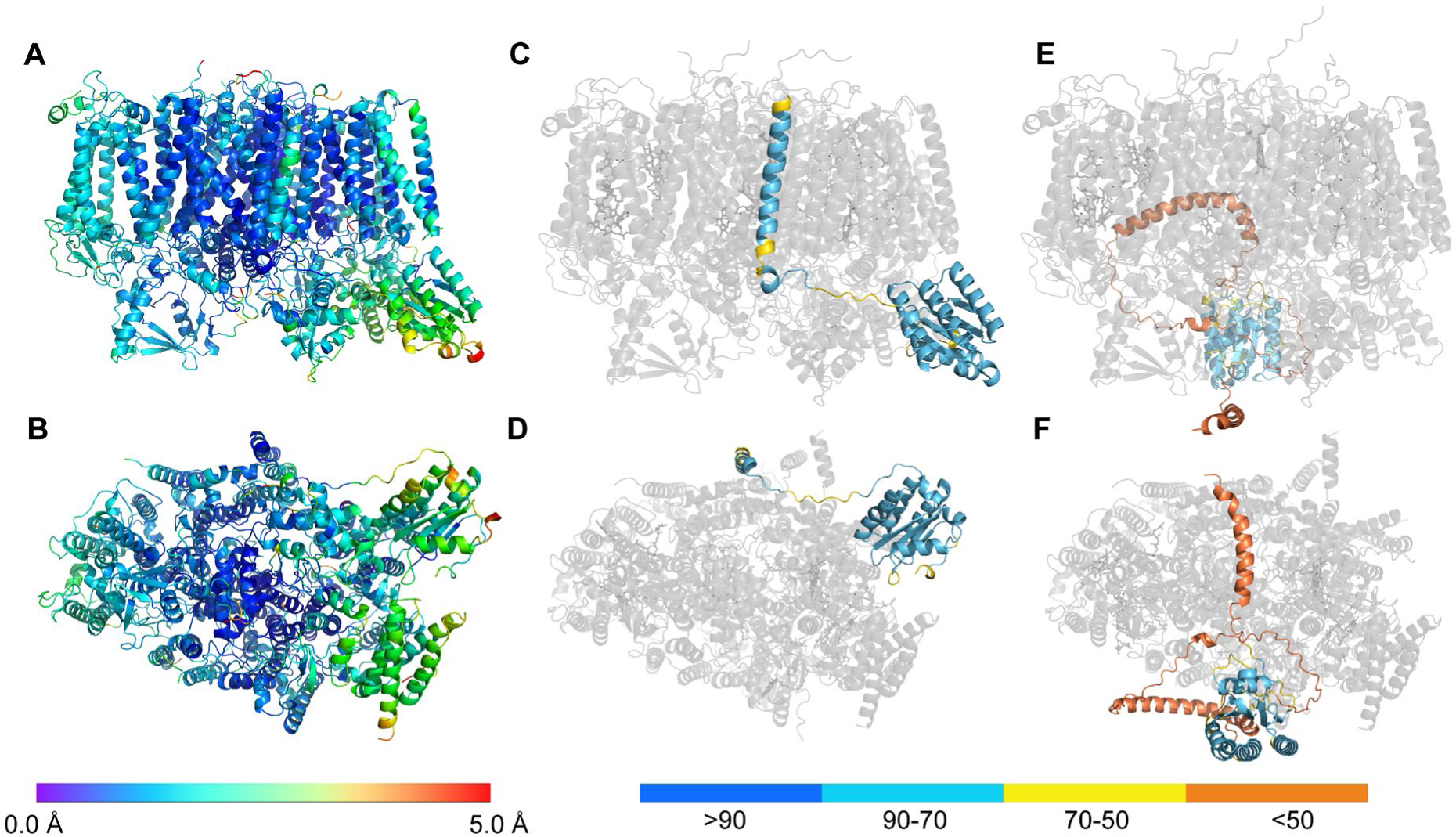
Structure predictions of Psb32 and the Psb32-like protein. Front (**A**) and bottom (**B**) view of the predicted Psb32-PSII colored according to the root mean square deviation of the Cɑ atom position compared to the cryo-EM model. Front (**C**) and bottom (**D**) view of the AlphaFold3^11^ predicted Psb32-PSII complex. Psb32 is colored according to the AlphaFold pIDDT score. The front (**E**) and bottom (**F**) view of the predicted complex with the Psb32-like protein (colored by plDDT) reveals a shifted binding position and a low pIDDT score for all but the central globular domain. Interestingly, AlphaFold predicts the TMH of Psb32 in all suggested models with high confidence at the same position as indicated by the cryo-EM density. Also, the Psb32 globular domain is predicted at the same position as indicated by the cryo-EM density mainly coordinated by PsbV and Psb27. Though the cryo-EM model was neither involved in training nor as template for the prediction the whole predicted Psb32-PSII complex and the cryo-EM model agree very well with a low overall Cα−RMSD of 1.30 Å (2563 atoms). This low RMSD implies that AlphaFold can accurately predict the Psb32-PSII assembly intermediate, since the cryo-EM structure was not included in the training set or used as template. However, the predicted TMH of the Psb32-like protein shows low confidence and lacks coordinating interactions. The globular domain of the Psb32-like protein is positioned in close proximity to PsbA and CP43, far from the location of Psb32. The prediction results of the Psb32-PSII and a tentative Psb32-like protein PSII assembly intermediate indicate that the Psb32-like protein cannot form a PSII assembly intermediate and therefore does not function as an assembly factor.

**Figure S11.**
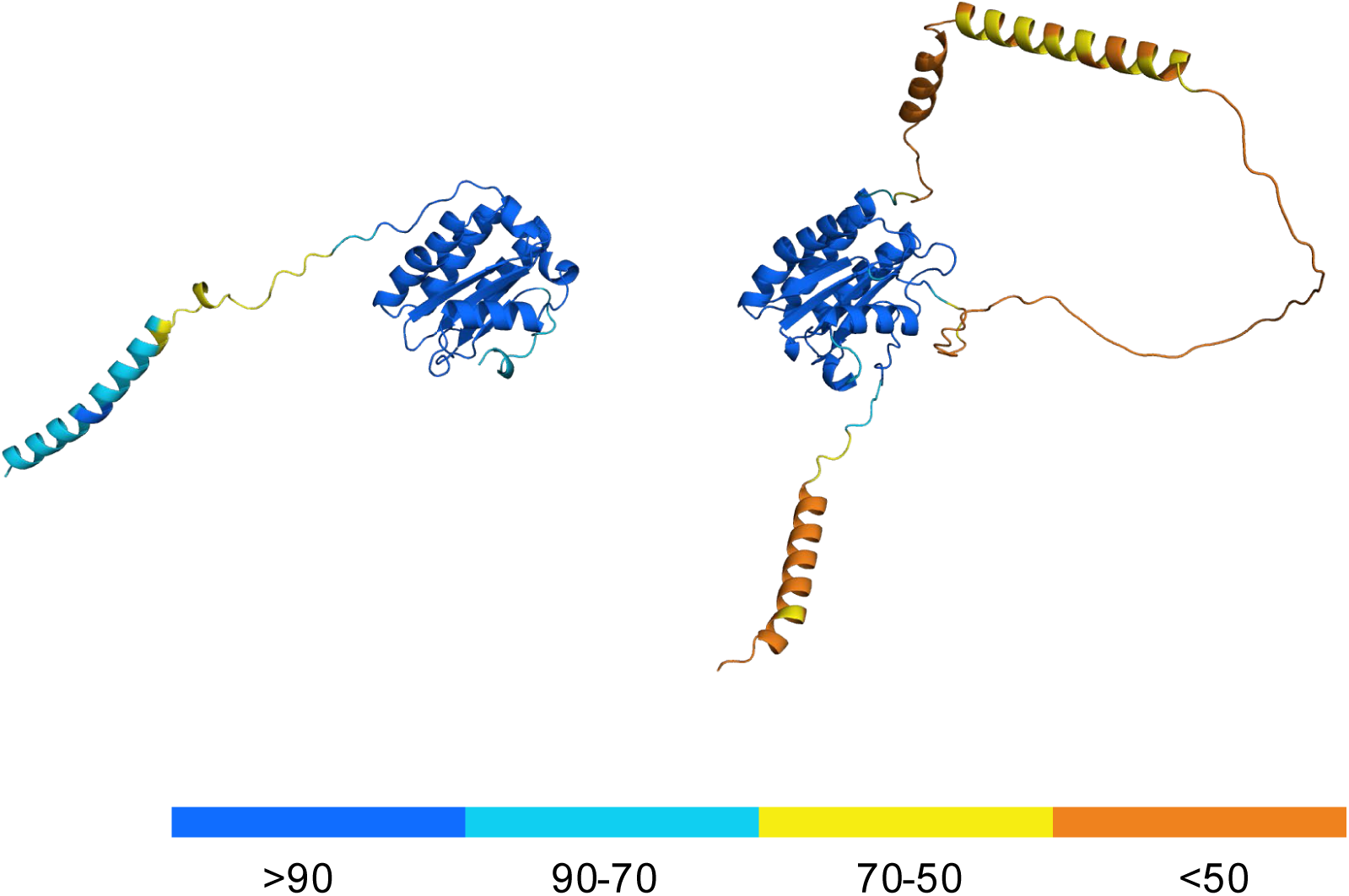
Comparison of predicted isolated Psb32 and Psb32-like protein structural models. The predicted Psb32 structure (**left,** colored according to their plDDT) shows high confidence in the globular domain of the protein and confidence in the C-terminal helix. The connecting unstructured linker shows partial low confidence. While the predicted Psb32-like protein structure (**right**) shows high confidence in the globular domain, its N- and C-terminal regions show low to very low confidence.

**Figure S12:**
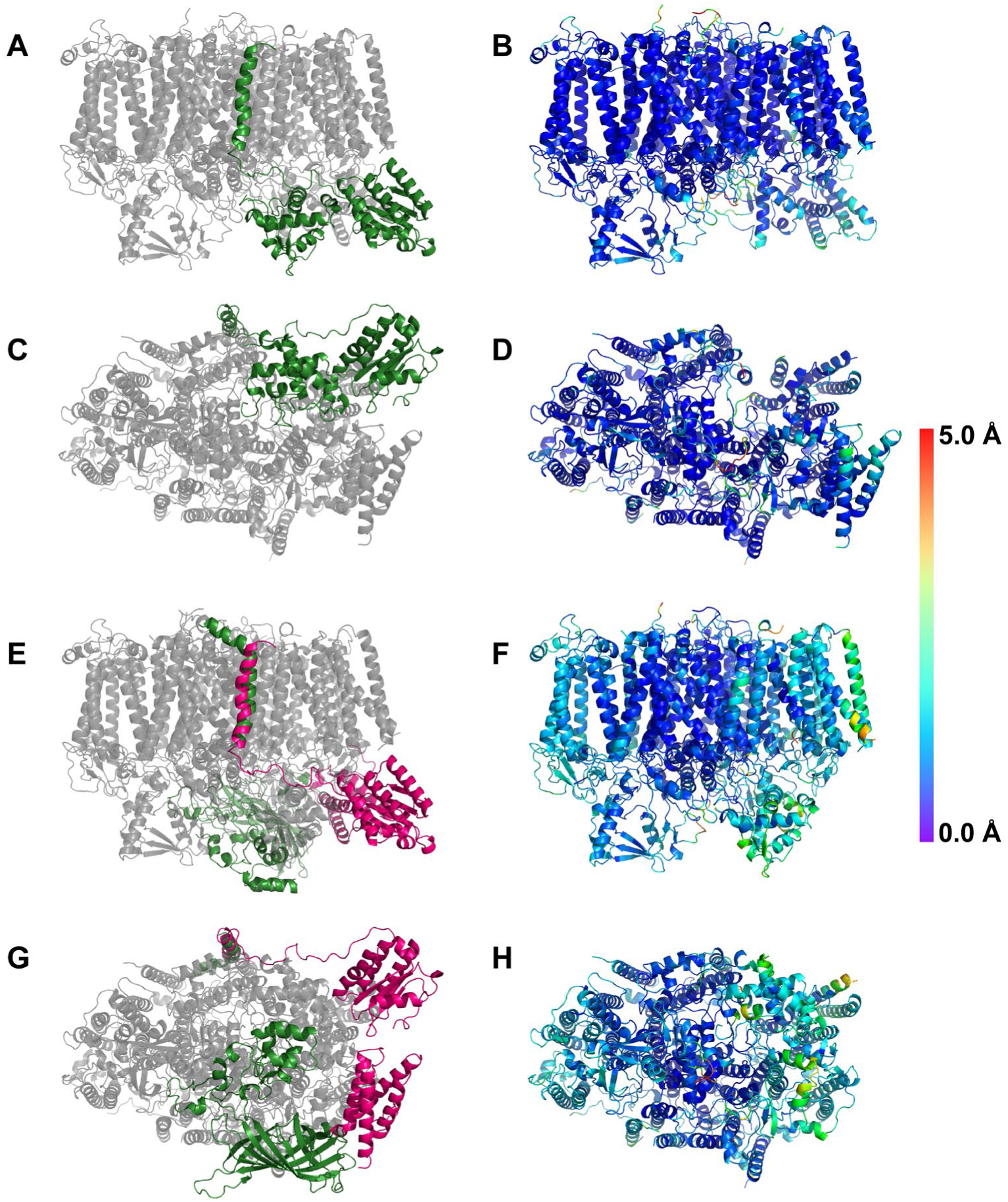
Changes within the PSII complex across the assembly. Front and bottom view of the leaving (red) and newly introduced (green) subunits upon switching from Psb27-PSII to Psb32-PSII (**A, C**) and switching from Psb32-PSII to the mature complex (**E, G**). Per-residue-colouring of the Cɑ-RMSD upon switching between Psb27-PSII and Psb32-PSII (**B, D**), and Psb32-PSII and the mature complex (**F, H**), respectively. The colouring ranges from 0 Å RMSD (deep blue) to 5 Å RMSD (red) between the respective models.

**Figure S13:**
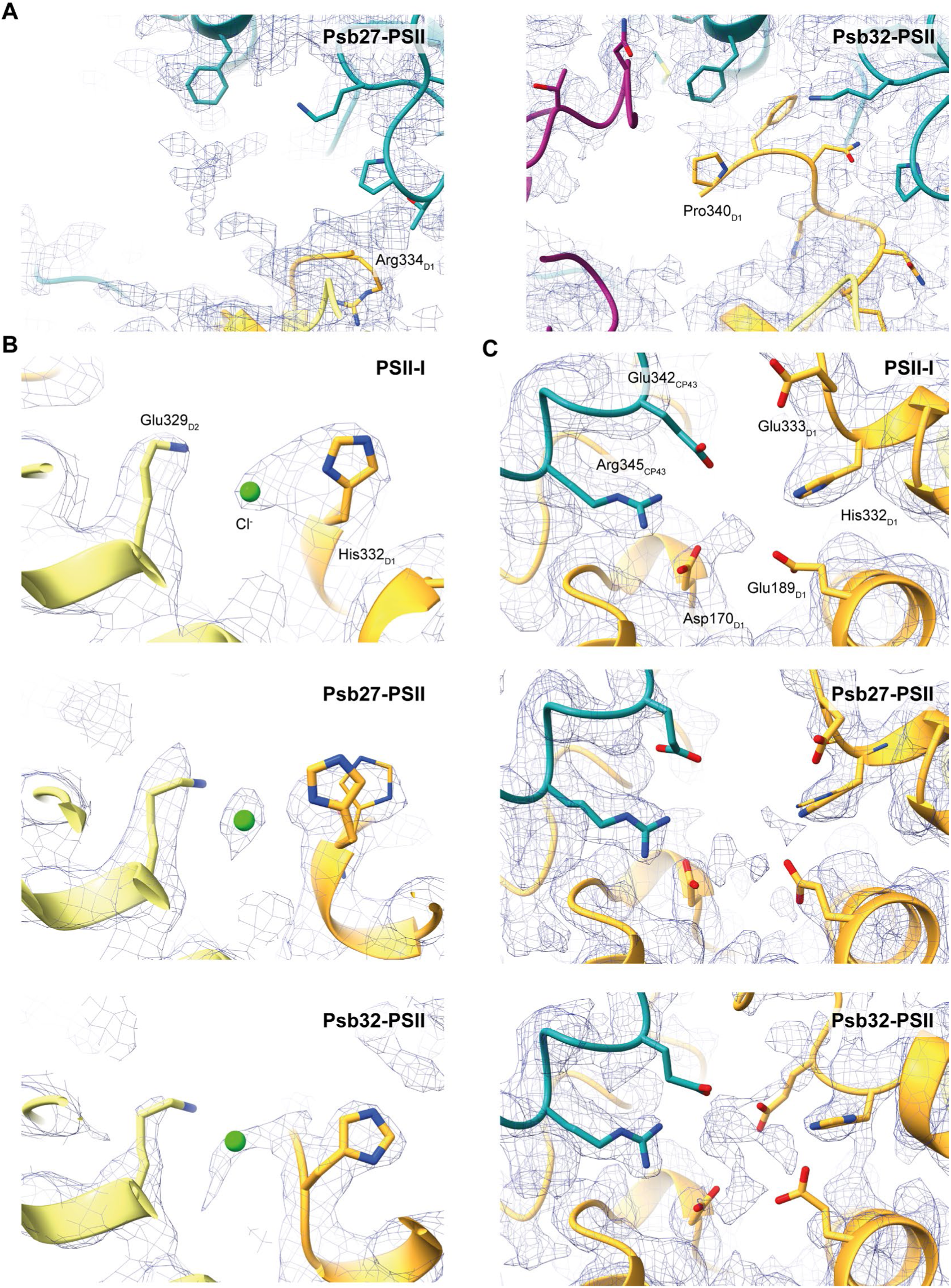
Cryo-EM density and model fitting around the OEC. **A)** Density fitting of Psb27-PSII D1 C-terminal loop (left) and Psb32-PSII D1 C-terminal loop (right). Maps are displayed at 2 σ. In Psb27-PSII, no defined density could be resolved beyond D1 Arg334 while in Psb32-PSII, no defined density could be resolved beyond D1 Pro340. **B)** Density fitting of His332 and chloride ion in PSII-I, Psb27-PSII and Psb32-PSII. **C)** Residues resolved around the OEC of PSII-I, Psb27-PSII and Psb32-PSII. For **B** and **C**, density maps of PSII-I, Psb27-PSII and Psb32-PSII are shown at 2.5 σ, 3.0 σ, and 3.5 σ, respectively.

